# Connexin26 mediates CO_2_-dependent regulation of breathing via glial cells of the medulla oblongata

**DOI:** 10.1101/2020.04.16.042440

**Authors:** Joseph Van de Wiel, Louise Meigh, Amol Bhandare, Jonathan Cook, Sarbjit Nijjar, Robert Huckstepp, Nicholas Dale

## Abstract

Breathing is highly sensitive to the PCO_2_ of arterial blood. Although CO_2_ is detected via the proxy of pH, CO_2_ acting directly via Cx26 may also contribute to the regulation of breathing. Here we exploit our knowledge of the structural motif of CO_2_-binding to Cx26 to devise a dominant negative subunit (Cx26^DN^) that removes the CO_2_-sensitivity from endogenously expressed wild type Cx26. Expression of Cx26^DN^ in glial cells of a circumscribed region of the medulla - the caudal parapyramidal area – reduced the adaptive change in tidal volume and minute ventilation by approximately 30% at 6% inspired CO_2_. As central chemosensors mediate about 70% of the total response to hypercapnia, CO_2_-sensing via Cx26 in the caudal parapyramidal area contributed about 45% of the centrally-mediated ventilatory response to CO_2_. Our data unequivocally links the direct sensing of CO_2_ to the chemosensory control of breathing and demonstrates that CO_2_-binding to Cx26 is a key transduction step in this fundamental process.

## Introduction

Breathing is a vital function that maintains the partial pressures of O_2_ and CO_2_ in arterial blood within the physiological limits. Chemosensory reflexes regulate the frequency and depth of breathing to ensure homeostatic control of blood gases (West, 2012). Historically, the ventral surface of the medulla oblongata has been recognized as an important location of central respiratory chemosensors (Loeschcke, 1982; Mitchell et al., 1963; Schlaefke et al., 1979; Schlaefke et al., 1970; Trouth et al., 1973b). Recent work has focused on two populations of neurons thought to contribute to the chemosensory control of breathing: those of the retrotrapezoid nucleus (RTN) (Guyenet, 2008; Kumar et al., 2015; Mulkey et al., 2006; Mulkey et al., 2004; Nattie et al., 1993) and the medullary raphé (Bernard et al., 1996; Brust et al., 2014; Ray et al., 2011; Richerson, 2004; Wang et al., 1998).

According to traditional consensus, CO_2_ is detected via the consequent change in pH, and pH is a sufficient stimulus for all adaptive changes in breathing in response to hypercapnia (Loeschcke, 1982). pH-sensitive K^+^ channels (TASKs and KIRs) are potential transducers. Although TASK-1 in the peripheral chemosensors of the carotid body (CB) contributes to overall pH/CO_2_ chemosensitivity (Trapp et al., 2008), TASK-1 does not appear to play a role in central pH/CO_2_ chemosensing (Guyenet et al., 2008). By contrast, TASK-2 may act as a central sensor of pH and contribute to adaptive changes in breathing (Kumar et al., 2015; Wang et al., 2013a). Recently a pH sensitive receptor, GPR4, has been linked to central chemosensitivity in the RTN. Complete deletion of this gene (from all tissues) greatly reduced the CO_2_ chemosensitivity in mice (Kumar et al., 2015). However, GPR4 is widely expressed in neurons including those of the medullary raphé, peripheral chemosensors, as well as the endothelium (Hosford et al., 2018). Moreover, systemic injection of a selective GPR4 antagonist modestly reduced the ventilatory response to CO_2_, but this same antagonist when administered centrally had no effect on the CO_2_-sensitivity of breathing (Hosford et al., 2018). A mechanism of pH-dependent release of ATP from ventral medullary glial cells may also contribute to the CO_2_-dependent regulation of breathing (Gourine et al., 2010).

There is considerable evidence that CO_2_ can have additional independent effects from pH on central respiratory chemosensors (Eldridge et al., 1985; Huckstepp and Dale, 2011; Shams, 1985). Cx26 is known to be present at the ventral medullary surface in the caudal and rostral chemosensory areas (Huckstepp et al., 2010b; Solomon et al., 2001). We have shown that CO_2_ directly binds to connexin26 (Cx26) hemichannels and causes them to open (Huckstepp et al., 2010a; Meigh et al., 2013). We have identified the critical amino acid residues that are necessary and sufficient for this process (Dospinescu et al., 2019; Meigh et al., 2013). Cx26 hemichannels can be gated by a number of stimuli. They are opened by voltage at potentials greater than -20 mV (Valdez Capuccino et al., 2019) and closed by acidification (Yu et al., 2007). Like all connexins, Cx26 hemichannels can be opened by removal of extracellular Ca^2+^ (Muller et al., 2002). However, the CO_2_-dependent opening of Cx26 hemichannels can occur in the absence of membrane depolarisation and at physiological levels of extracellular Ca^2+^ (de Wolf et al., 2017; Dospinescu et al., 2019; Huckstepp et al., 2010a; Meigh et al., 2013).

This direct gating of Cx26 is an important new mechanism that underlies CO_2_-dependent ATP release (Gourine et al., 2005; Huckstepp et al., 2010a; Huckstepp et al., 2010b; Wenker et al., 2012) and provides a potential mechanism for the direct action of CO_2_ on breathing. Nevertheless, the role of direct sensing of CO_2_ in the regulation of breathing remains uncertain because genetic evidence linking Cx26 to the control of breathing has been lacking, and the cells that could mediate direct CO_2_ sensing via Cx26 have not been identified.

In this study we have addressed both of these issues by exploiting our knowledge of the binding of CO_2_ to Cx26 to devise a dominant negative subunit (Cx26^DN^) that removes CO_2_- sensitivity from endogenous wild type Cx26 hemichannels. By using a lentiviral construct to drive expression of Cx26^DN^ in glial cells of the ventral medulla, we have obtained evidence that links CO_2_-dependent modulation of Cx26 in glial cells present in a small circumscribed area of the ventral medulla to the adaptive control of breathing.

## Results

### Rationale for the design of Cx26^DN^

CO_2_ causes Cx26 hemichannels to open via carbamylation of K125 and subsequent formation of a salt bridge between the carbamylated lysine side chain and R104 of the neighbouring subunit (Figure 1a) (Meigh et al., 2013). As Cx26 hemichannels are hexameric there are potentially 6 binding sites for CO_2_ suggesting the potential for highly cooperative binding of CO_2_. Indeed, Cx26 is steeply sensitive to changes in PCO_2_ around its physiological level of about 40 mmHg (Figure 1b) (Huckstepp et al., 2010a). We reasoned that introducing two mutations: K125R, to prevent CO_2_-dependent carbamylation, and R104A to prevent salt bridge formation (“carbamate bridges”) to a neighbouring carbamylated subunit, should produce a dominant negative subunit (Figure 1c). If such a subunit coassembled into the Cx26 hexamer, it would likely have a dominant negative action as it would remove the capacity to form at least two out of the 6 possible carbamate bridges (Figure 1c).

### Cx26^DN^ coassembles with Cx26^WT^

To determine whether the dominant negative subunit will effectively coassemble into a hexamer with the wild type subunit, we used acceptor depletion FRET (Figure 2). We exploited the Clover-mRuby2 FRET pair (Lam et al., 2012) by tagging Cx26^WT^ and Cx26^DN^ with Clover (donor) and mRuby2 (acceptor). When mRuby2 was bleached in HeLa cells coexpressing either Cx26^WT^-Clover and Cx26^WT^-mRuby2 or Cx26^WT^-Clover and Cx26^DN^-mRuby2, the fluorescence of the Clover was enhanced where these two fluorophores were colocalised (Figure 2). The FRET efficiency for Cx26^WT^-Clover and Cx26^WT^-mRuby2 or Cx26^WT^-Clover and Cx26^DN^-mRuby2 was very similar (Figure 3). This suggests that the Cx26^DN^ subunit interacts closely with Cx26^WT^. In principle, this could be because a homomeric hexamer of Cx26^WT^ was sufficiently close to a homomeric hexamer of Cx26^DN^ by random association in the membrane rather than assembly of a heteromeric hexamers comprised of both types of subunits. We therefore also looked at FRET interactions between Cx43^WT^ and Cx26^WT^. These two connexin subunits do not form heteromeric hexamers, but homomeric hexamers of the two types could come close to each other in the plasma membrane by random association. We found that the FRET efficiency of this interaction was much lower than that of Cx26^WT^-Cx26^WT^ or Cx26^WT^-Cx26^DN^ (Figures 2,3). This suggests that Cx26^DN^ does indeed coassemble into a hexamer with Cx26^WT^. This hypothesis gains further support from our analysis that shows that the FRET efficiency for Cx26^WT^-Cx26^DN^ is negatively correlated with the Donor to Acceptor (D-A) ratio (Supplementary Figure 1) which is indicative of assembly into heteromeric hemichannels (Di et al., 2005). By contrast the FRET efficiency for Cx43^WT^-Cx26^WT^ shows no such correlation with the D-A ratio (Supplementary Figure 1). There is a positive correlation of FRET efficiency with Acceptor level which may indicate some additional random association of homomeric connexons comprised of Cx26^WT^ and Cx26^DN^ (Di et al., 2005) (Supplementary Figure 1). Nevertheless, our data strongly indicate that the Cx26^DN^ subunit effectively coassembles with Cx26^WT^ subunits to form heteromeric hemichannels.

**Figure 1.**
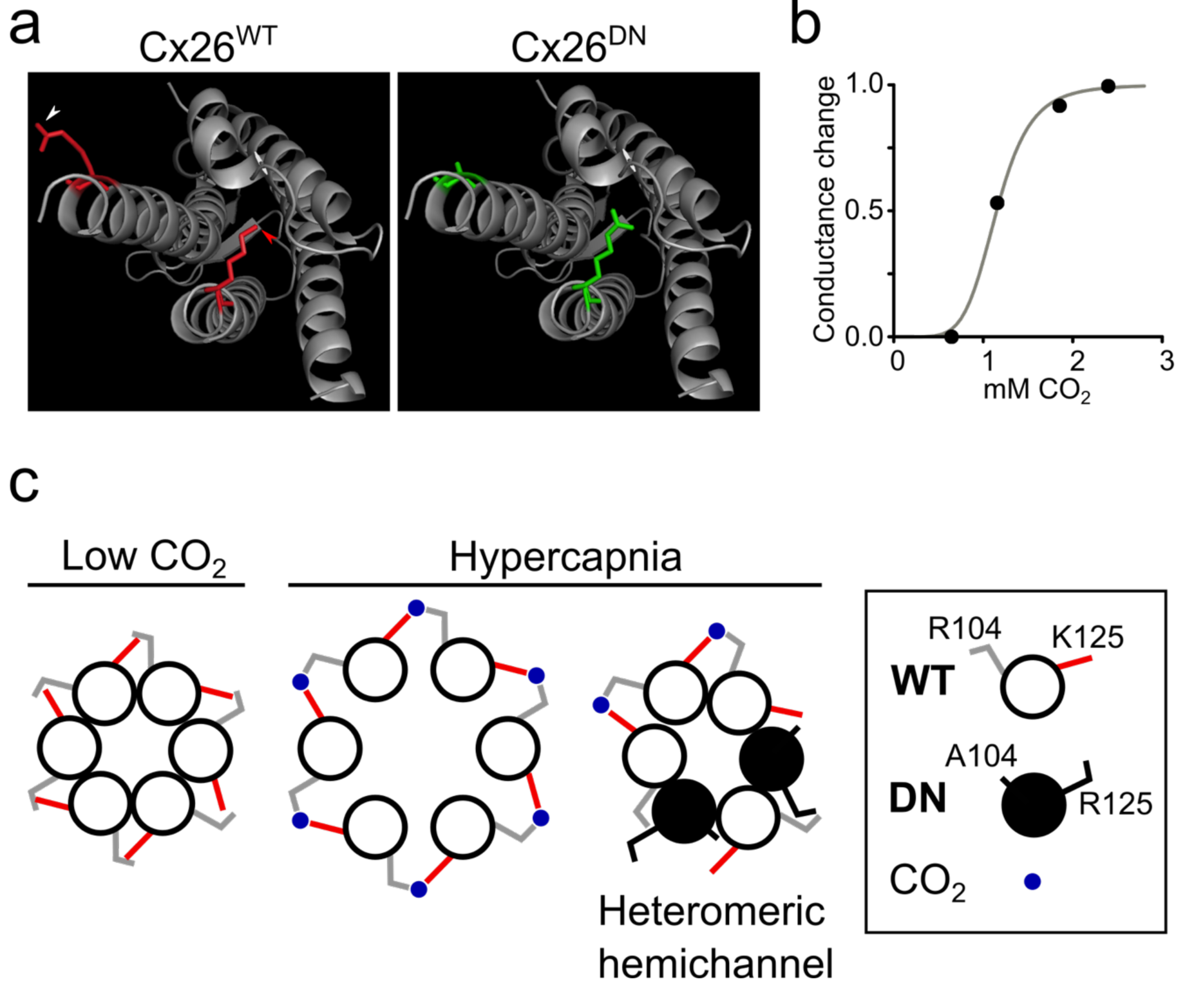
Rationale for the design of a dominant negative Cx26 subunit (Cx26^DN^) to inhibit CO_2_-mediated hemichannel opening. a) Ribbon diagram, showing a single Connexin 26 subunit, highlighting the two amino acid residues vital for CO_2_ sensing - R104 (white arrowhead) and K125 (red arrowhead, Cx26^WT^, left). In the Cx26^DN^ mutant these two residues are mutated - R104A and K125R (Cx26^DN^, right). This will prevent carbamylation of the subunit and formation of a salt “carbamate” bridge with adjacent connexin subunit in the hexamer. b) CO_2_-dependent whole-cell clamp conductance changes from HeLa cells stably expressing Cx26^WT^ (data from (Huckstepp et al., 2010a)). The grey line is drawn with a Hill coefficient of 6, suggesting that hemichannel opening to CO_2_ is highly cooperative. c) Hypothesised coassembly of Cx26^WT^ and Cx26^DN^ into heteromeric hemichannels that will be insensitive to CO_2_ because insufficient carbamate bridge formation will occur to induced channel opening.

**Figure 2.**
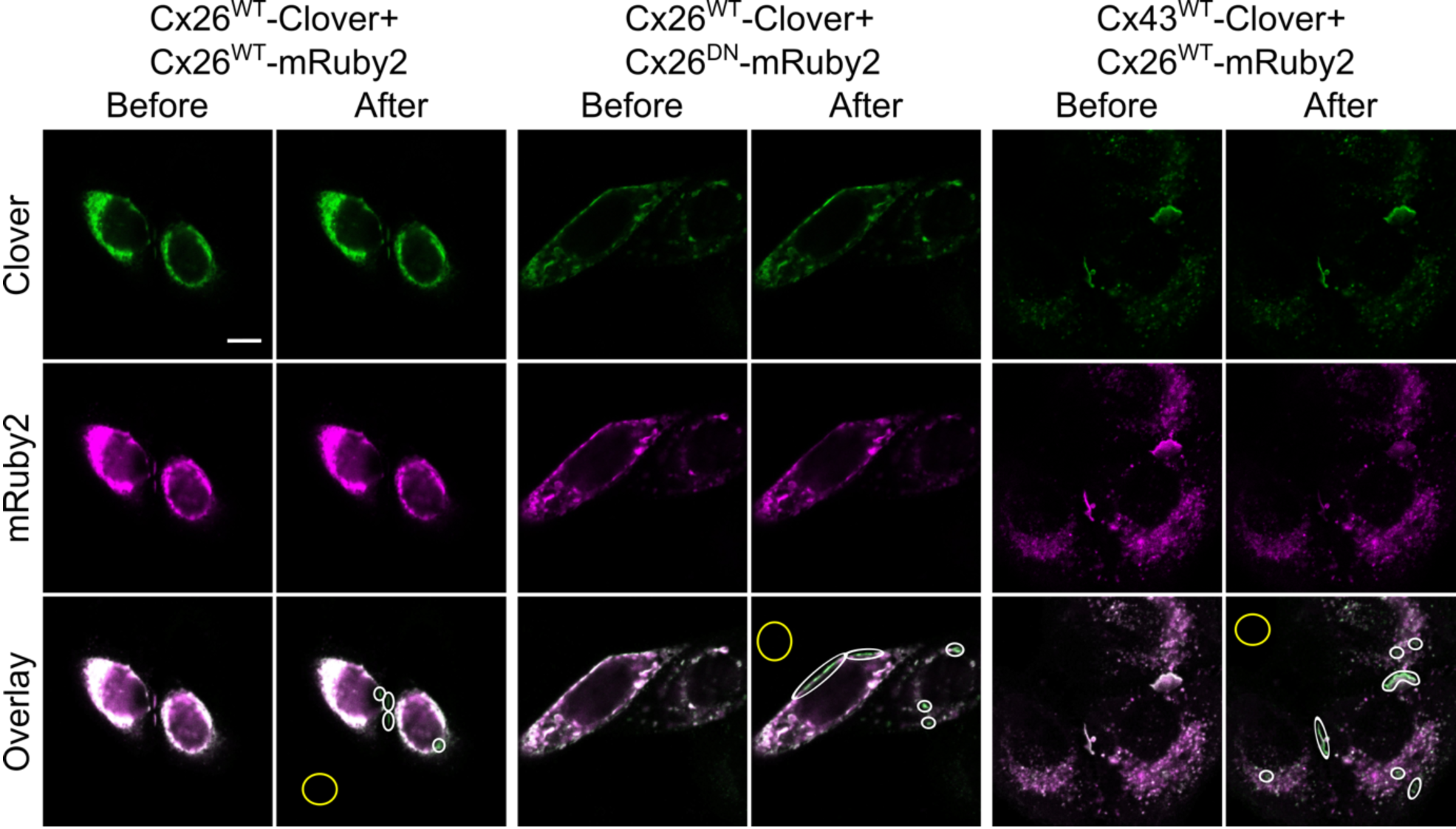
FRET signal between connexin variants. Example images of acceptor depletion Fö rster Resonance Energy Transfer (ad-FRET) experiments. HeLa cells were co-transfected with equal amounts of DNA transcripts (to express connexin-fluorophore constructs) and PFA fixed after 48 hrs later. Two channels were recorded: 495-545nm (Clover, green) and 650-700nm (mRuby2, magenta), and images were acquired sequentially with 458nm and 561nm argon lasers (respectively). Photobleaching was performed using the 561nm laser for 80 frames at 100% power, targeting ROIs. Three combinations of connexin-Clover (donor, green) and connexin-mRuby2 (acceptor, red) are shown before and after bleaching. Colocalisation is shown in white in the overlay images, along with bleached areas (white ovals) and background references (yellow ovals). Following acceptor bleaching, ROIs in Cx26^WT^-Clover + Cx26^WT^-mRuby2 and Cx26^WT^-Clover + Cx26^DN^-mRuby2 samples show enhanced fluorescence intensity of donor (Clover, green) and reduced fluorescence intensity of acceptor (mRuby2, magenta). ROIs in Cx43^WT^-Clover + Cx26^WT^-mRuby2 samples show almost no change in fluorescence intensity of donor (Clover, green) and reduced fluorescence intensity of acceptor (mRuby2, magenta). Colocalisation measurements decrease mainly due to photobleaching of mRuby2, and subsequent elimination of its fluorescence. All co-transfection combinations showed statistically significant colocalisation at the highest resolution for the images. Colocalisation threshold calculations were carried out in ImageJ using the Costes method; 100 iterations, omitting zero-zero pixels in threshold calculation. Statistical significance was calculated for entire images and individual ROIs, with no difference between either calculation (p-value=1 for all images, with a significance threshold of p>0.95). Scale bar, 10 μm.

**Figure 3.**
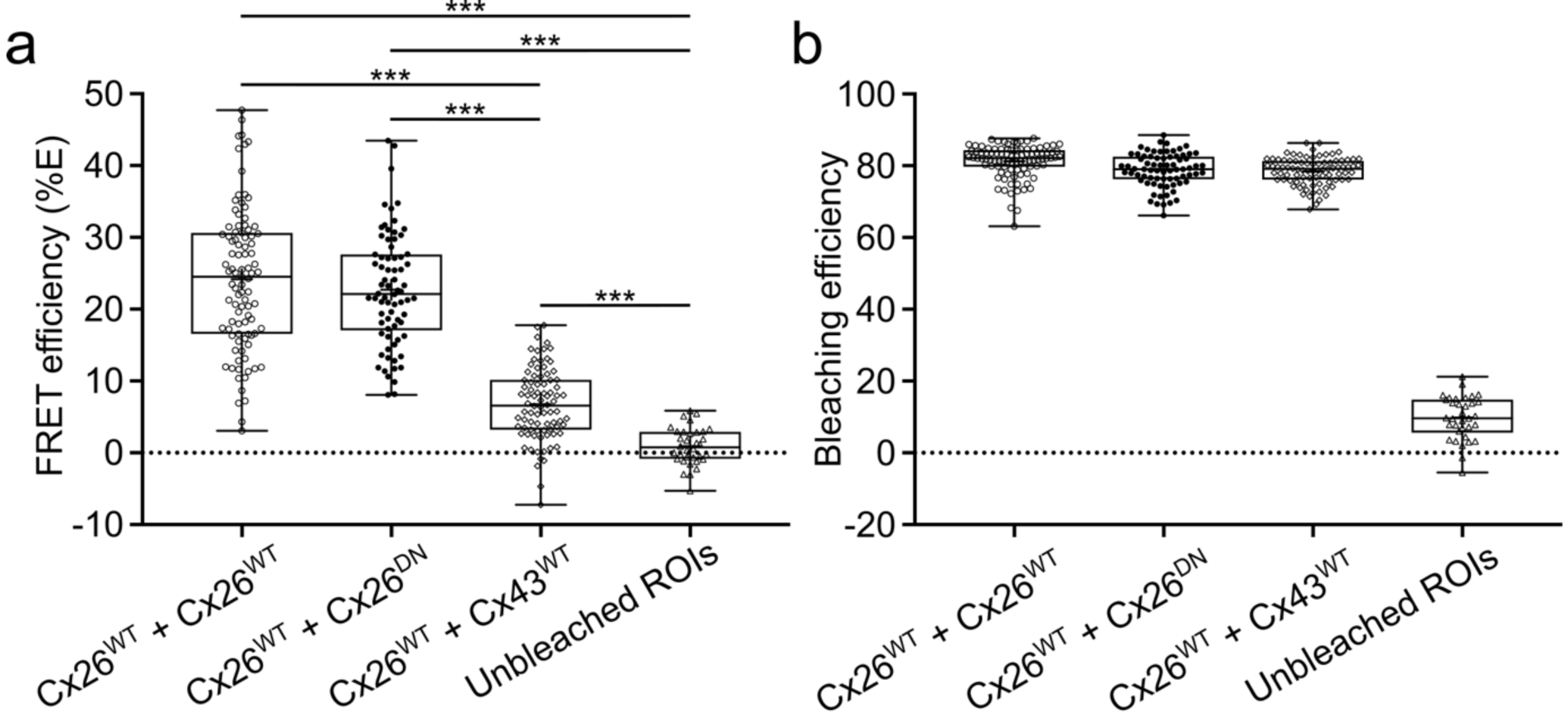
FRET efficiency of coexpressed connexin variants. a) Box and whisker plots showing the difference in FRET efficiency (%E) across different connexin co-expression connexin samples. FRET efficiency was calculated from background-adjusted ROIs as: %E = 100 X (clover_Post_ - clover_Pre_)/clover_Post_. One-way ANOVA was carried out in SPSS. Post-hoc testing revealed all individual comparisons to be significant at p<0.001 (***), with the exception of the comparison between Cx26^WT^ + Cx26^WT^ and Cx26^WT^ + Cx26^DN^ datasets which were not significantly different. Each dot represents a different ROI. b) Box and whisker plot showing mRuby2 bleaching efficiency during the acceptor depletion step. Whilst a small amount of bleaching occurred in untargeted regions (presumably due to light scattering and/or reflection), targeted ROI’s received drastically greater bleaching. Importantly all targeted regions showed highly similar bleaching efficiencies. Bleaching efficiency was calculated from background-adjusted ROIs as: Bleaching = (1 – (mRuby2_Post_/mRuby2_pre_)). Boxes show the interquartile range, the median is indicated by the horizontal line within the box, and the mean is indicated by the cross within the box. Range bars show minimum and maximum values.

### Cx26^DN^ has a dominant negative action on the CO_2_-sensitivity of Cx26^WT^

We have previously shown that hemichannels comprised of Cx26^K125R^ and Cx26^R104A^ are insensitive to CO_2_ (Meigh et al., 2013). We therefore confirmed that Cx26^DN^ hemichannels, which carry both of these mutations, are also insensitive to CO_2_ (Supplementary Figure 2). This important observation shows that if Cx26^DN^ were to form homomeric hemichannels *in vivo* they would have no impact on the CO_2_-sensitivity of cells expressing them.

To assess the capacity of Cx26^DN^ to act as a dominant negative construct with respect to CO_2_ sensitivity, we transfected this subunit into HeLa cells that stably expressed Cx26^WT^. We then used a dye loading assay to assess how the sensitivity of these HeLa cells to CO_2_ changed with time. Four days after transfection, the HeLa cells that coexpressed Cx26^DN^ with Cx26^WT^were as sensitive to CO_2_ as those HeLa cells that only expressed Cx26^WT^ (Figure 4). However, 6 days after transfection, the HeLa cells that coexpressed Cx26^DN^ were insensitive to CO_2_ (Figure 4). We conclude that Cx26^DN^ has a dominant negative effect on the CO_2_ sensitivity of Cx26^WT^ and exerts this effect by coassembling into heteromeric hexamers with Cx26^WT^ in the manner hypothesised in Figure 1.

**Figure 4.**
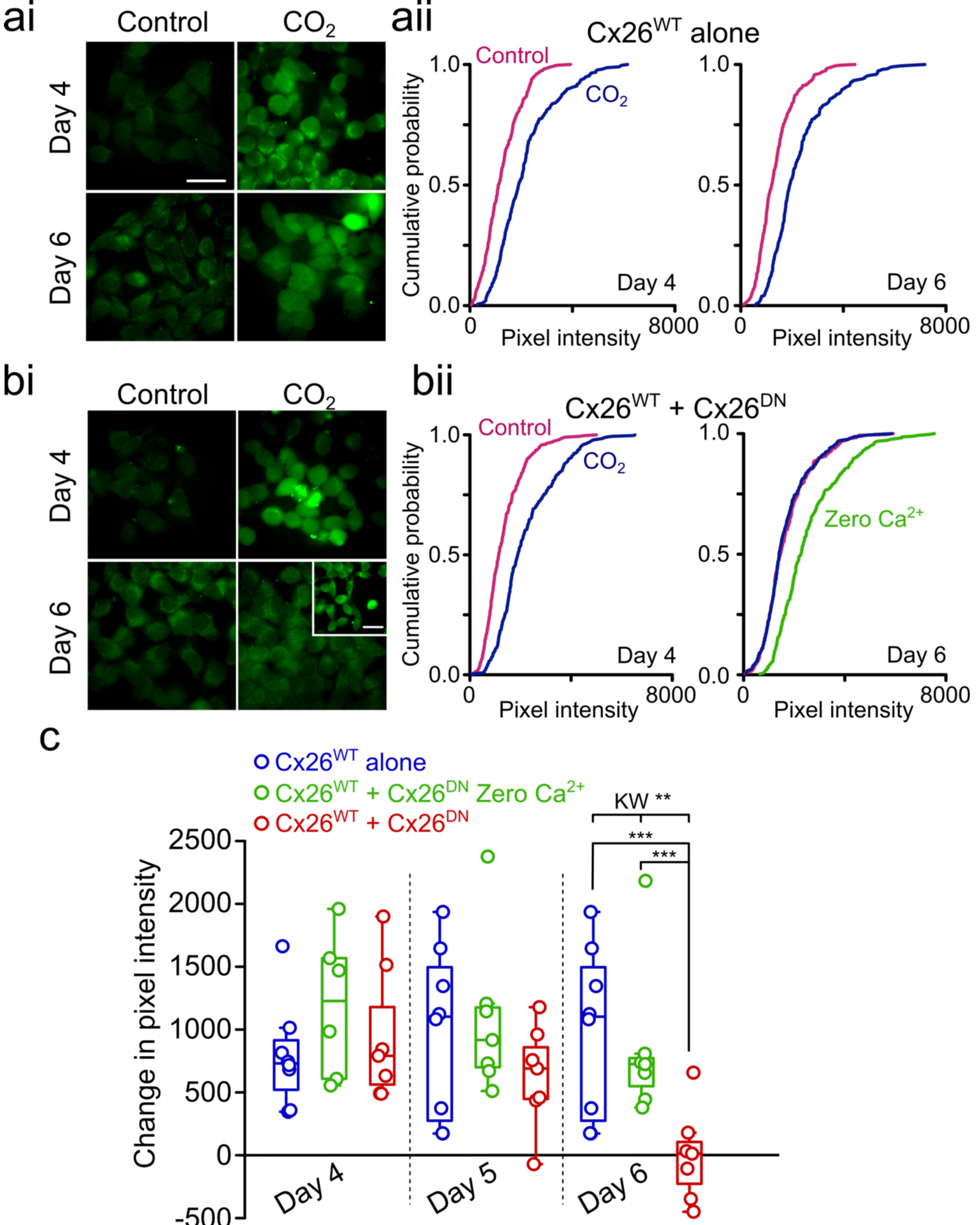
Cx26^DN^ removes CO_2_-sensitivity in HeLa cells stably expressing Cx26^WT^. Dye-loading under 35mmHg PCO_2_ (control) or 55mmHg PCO_2_ (hypercapnic) conditions revealed how the CO_2_ sensitivity of HeLa cells stably expressing Cx26 (Cx26-HeLa cells) changes over time after transfection with Cx26^DN^. ai, bi) Representative dye-loading images of Cx26-HeLa cells at day 4 and day 6 days after cells were either untreated (ai) or transfected with Cx26^DN^ (bi). In (bi) the inset represents a Zero Ca^2+^ control to demonstrate the presence of functional hemichannels even when the HeLa cells showed no CO_2_ dependent dye loading. aii, bii) Cumulative probability distributions comparing mean pixel intensity for each condition at day 4 and day 6 (untreated Cx26-HeLa cells, b; Cx26-HeLa cells transfected with Cx26^DN^. n>40 cells per treatment repeat, with 5 independent repeats for each treatment. The cumulative distributions show every data point (cell fluorescence intensity measurement). c) Median change in pixel intensity caused by 55 mmHg PCO_2_ and Zero Ca^2+^ from baseline (35mmHg) over days 4, 5, and 6 post-transfection. At Day 6, Median pixel intensities (from 7 independent repeats) were compared using the Kruskal Wallace ANOVA (p=0.007), and post-hoc with the Mann-Whitney U test (Cx26^WT^ vs Cx26^WT^+Cx26^DN^, p=0.003; Cx26^WT^+Cx26^DN^ Zero Ca^2+^ vs Cx26^WT^+Cx26^DN^, p=0.002). Each circle represents one independent replication (independent transfections and cell cultures). Scale bars, 40μm.

### In vivo expression of Cx26^DN^ blunts the CO_2_ sensitivity of breathing

The Cx26^DN^ subunit has the potential to be a genetic tool that could remove CO_2_ sensitivity from Cx26 and thus probe this aspect of Cx26 function without deleting the Cx26 gene. This has the advantage of leaving other signalling roles of Cx26 intact, and also of linking any CO_2_-dependent physiological functions to the CO_2_binding site of Cx26 itself. We therefore created a lentiviral construct that contained either Cx26^DN^ or Cx26^WT^ (a control for the Cx26^DN^ construct) under the control of a bidirectional cell-specific GFAP promoter (Liu et al., 2008) (Supplementary Figure 3a). Rather than tagging a fluorescent protein to the C-terminus of the Cx26 variants, we expressed Clover behind an IRES so that the cells that expressed the Cx26 variants could be identified and quantified (Supplementary Figure 3a). Stereotaxic injection of the lentiviral construct into the ventral medulla oblongata, showed that lentiviral construct drove expression of the Cx26 variants selectively in GFAP+ cells (Supplementary Figure 3b).

We used bilateral stereotaxic injections of the lentiviral constructs into the medulla oblongata to assess the effect of Cx26^DN^ on the CO_2_ sensitivity of breathing by means of whole-body plethysmography (Supplementary Figure 4). Pilot experiments suggested that transduction of glial cells at the ventral medullary surface in a region ventral and medial to the lateral reticular nucleus reduced the adaptive changes in breathing to hypercapnia 3 weeks after injection of the virus (Supplementary Figure 5). From this pilot work, we designed a study to achieve a statistical power of 0.8 at a significance level of 0.05 in which we injected 12 mice with Cx26^WT^ and 12 mice with Cx26^DN^ and followed how the CO_2_ sensitivity of breathing changed with time following the lentiviral injection (Figure 5a). Two weeks after viral transduction the change in tidal volume to 6 % CO_2_ in the Cx26^DN^-injected mice was less than that of the Cx26^WT^-injected mice (two-way mixed effects ANOVA, p=0.008; post hoc one tailed t-test, p=0.002). The change in minute ventilation was also less in the Cx26^DN^-injected mice compared to the Cx26^WT^-injected mice (two-way mixed effects ANOVA, p=0.015; post hoc one tailed t-test, p=0.007). There was no readily discernible difference in the changes in respiratory frequency to CO_2_ between the Cx26^WT^ and Cx26^DN^-injected mice. Three weeks following the viral injection these differences had disappeared, presumably due to some compensatory mechanism within the respiratory networks (Forster, 2018; Johnson and Mitchell, 2013). Following the pilot study, we were able to place our injection more medially into the caudal parapyramidal area. More accurately targeting the injection of virus particles to the correct area, likely led to the quicker onset of the phenotype, as transfection of the relevant cells would thus have been more efficient and rapid.

**Figure 5.**
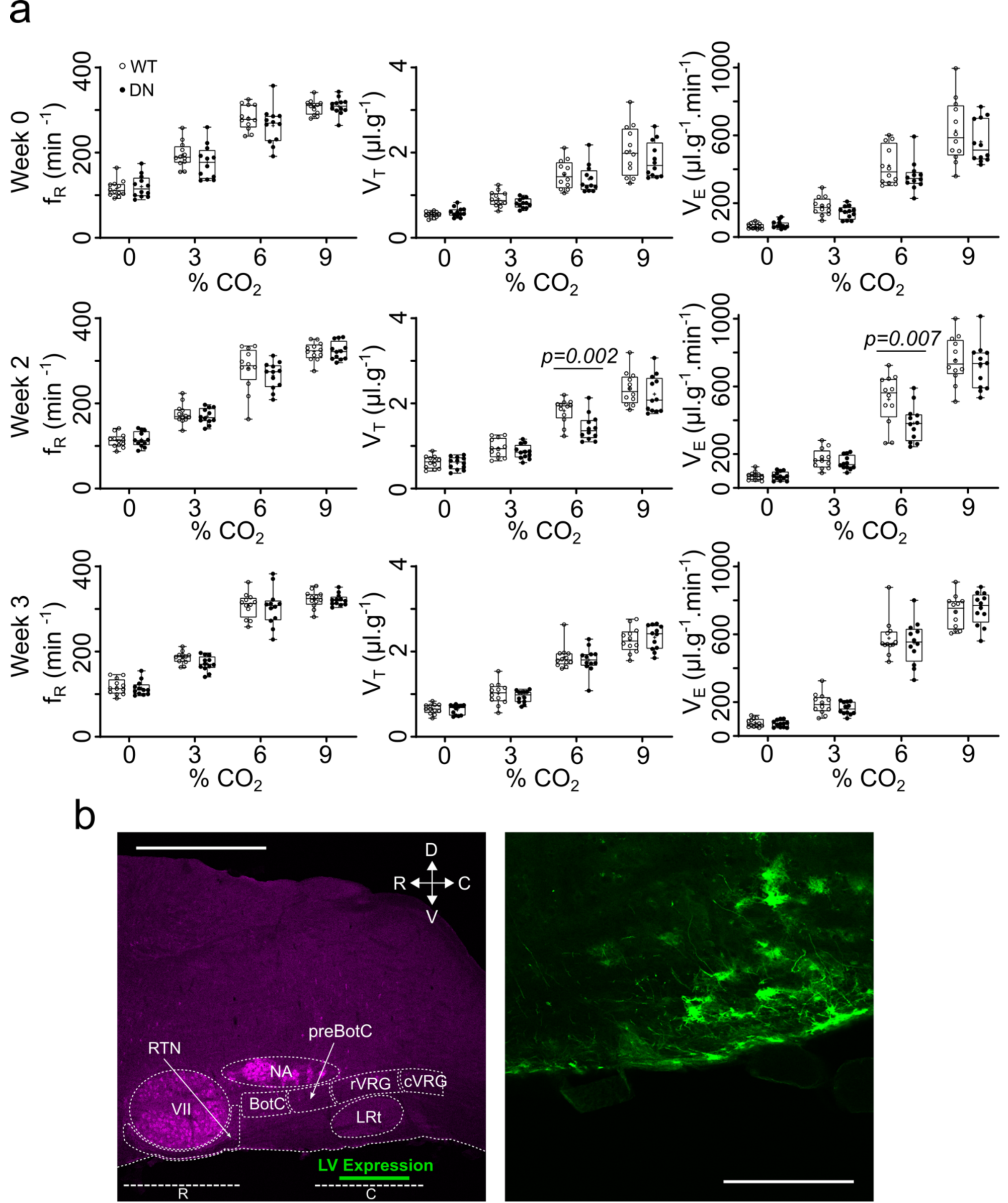
Connexin26-mediated hypercapnic breathing response in conscious mice. a) Mice aged 11-14 weeks were bi-laterally injected with lentivirus (LV) at the ventral medullary surface (VMS) to introduce either the Cx26^DN^ or Cx26^WT^ gene under the control of a GFAP promoter into genomic DNA. Whole-body plethysmography measurements of frequency (f_R_), tidal volume (V_T_), and minute ventilation (V_E_) were recorded for each mouse at 0%, 3%, 6%, and 9% CO_2_, before (week 0) and after (weeks 2 and 3) LV transduction. Two weeks after transduction there was a difference (two-way mixed ANOVA followed by post-hoc t-test, p values given for post hoc comparisons) in adaptive changes in tidal volume to 6% CO_2_ when comparing mice expressing Cx26^WT^ (empty circles, n = 12) and mice expressing Cx26^DN^ (black circles, n = 12). The median is indicated as a horizontal line within the box, and the mean is represented by a cross within the box. Range bars show minimum and maximum values. b) Location of GFAP:Cx26 LV construct expression (green) in the sagittal plane - scale bars, 1 mm (left), 200 µm (right). R, rostral chemosensitive site; C, caudal chemosensitive site; NA, nucleus ambiguous; VII, facial nucleus; preBot, preBötzinger Complex; Bot, Bötzinger Complex; rVRG, rostral ventral respiratory group; RTN, retrotrapezoid nucleus; cVRG, caudal ventral respiratory group; LRt, lateral reticular nucleus.

*Post hoc* tissue staining confirmed the location of the transduced glial cells, and highlighted cells at the very surface of the ventral medulla that had long processes projecting rostrally and dorsally into the parenchyma of the medulla (Figure 5b). The difference between Cx26^WT^ and Cx26^DN^ transduced mice could not be attributed to the increased expression of Cx26^WT^ and hence CO_2_ sensitivity, as sham operated mice (performed as part of the same study) showed no difference in CO_2_ sensitivity from the Cx26^WT^ mice (Supplementary Figure 7). In both the pilot experiment and main experiment, the effect of Cx26^DN^ was to alter the relationship of tidal volume and minute ventilation versus inspired CO_2_: specifically, Cx26^DN^ reduced the increase in both of these parameters that occurs at 6% inspired CO_2_ by ∼30% compared to the control (Cx26^WT^, see Supplementary Figures 5 and 6, and Supplementary Tables 1 and 2).

Expression of Cx26^DN^ in glial cells in this location ventral and medial to the lateral reticular nucleus appeared to be uniquely able to alter the CO_2_ sensitivity of breathing. Transduction of glial cells in the RTN (Supplementary Figure 8) or even more caudally in the medulla (Supplementary Figure 9) had no effect on the CO_2_ sensitivity of breathing. We therefore conclude that Cx26 in a circumscribed population of GFAP+ cells ventral and medial to the lateral reticular nucleus contributes to the CO_2_-dependent regulation of breathing. This is the first mechanistic evidence that shows a direct effect of CO_2_ on breathing and links the structural biology of CO_2_ binding to Cx26 to the regulation of breathing. While the lentiviral construct transduced typical astrocytes (e.g. Figure 5b) it consistently transduced glial cells that had a soma at the very ventral edge of the medulla. These glial cells were unusual in that they had very long processes, which extended rostrally (Figure 5a, Figure 6) and also medially (Figure 6).

**Figure 6.**
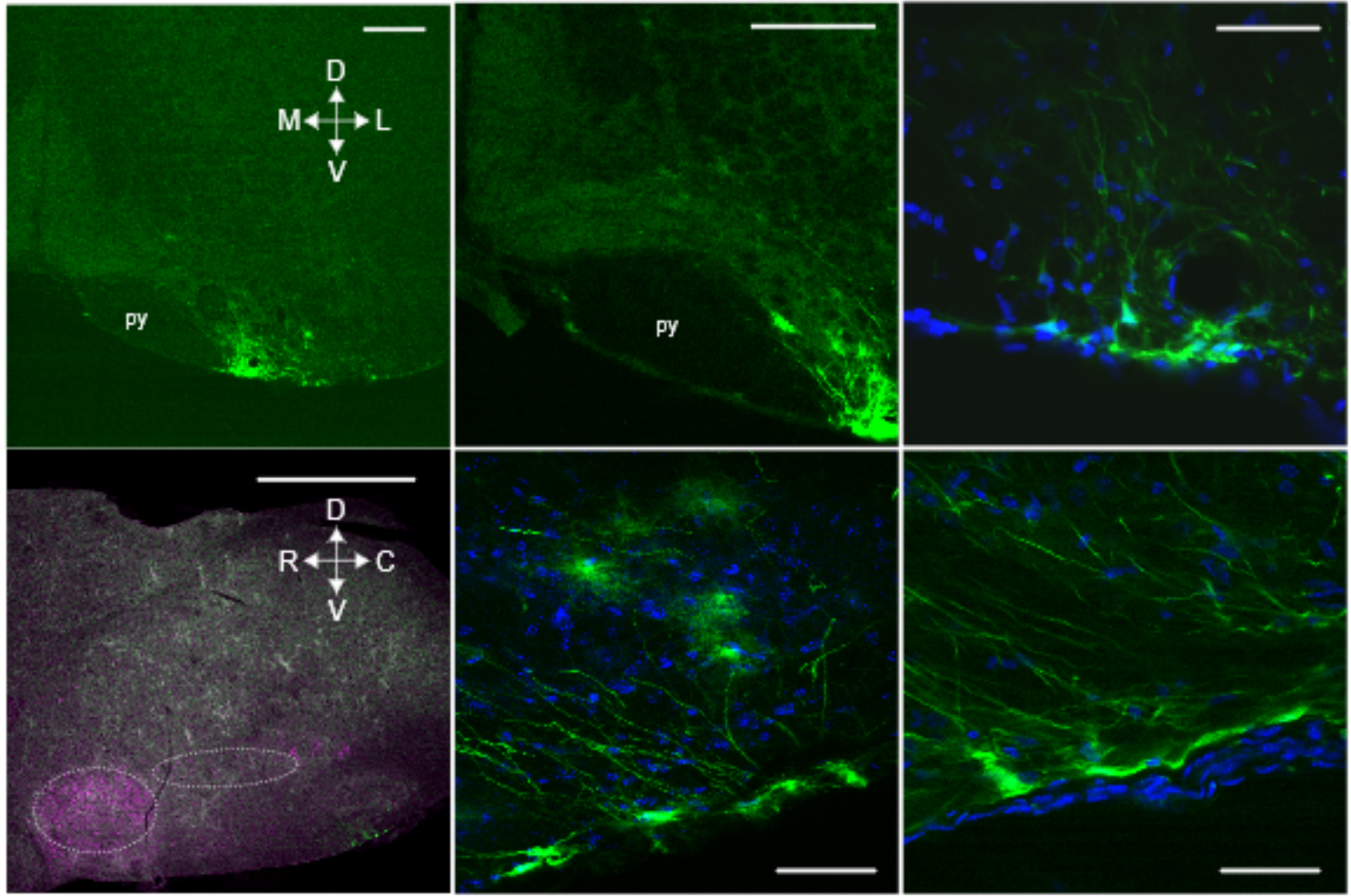
The putative chemosensory cells extend processes dorsal, medial, and rostral. Top 3 images: Coronal sections. Bottom 3 images: parasagittal sections showing the location of the transduced cells (left) and examples of the cells at higher magnification (middle and right). Cells expressing GFAP:Cx26:Clover (green) at the ventral medullary surface have a morphology unlike that of astrocytes, with a cell body at the very margin of the ventral surface and long processes that extend deep into the brain in the direction of respiratory nuclei. The ventrolateral respiratory column lies caudal to the VII nucleus and ventral to the nucleus ambiguous (dashed ovals). Choline acetyltransferase staining (magenta). py, pyramids. Scale bars: bottom-left, 1 mm; top-left and top-middle, 200 µm; top-right and bottom-middle and bottom-right, 50 µm.

## Discussion

By devising a dominant negative subunit Cx26^DN^, which coassembles with endogenously expressed Cx26^WT^ to remove CO_2_ sensitivity from the resulting heteromeric hemichannels, we have demonstrated a clear link between Cx26-mediated CO_2_-sensing and the regulation of breathing. Our study also links the structural motif of CO_2_ binding in Cx26 - the “carbamylation motif” – to the CO_2_-dependent regulation of breathing. CO_2_ carbamylation, happens spontaneously, and was originally described as the basis of the CO_2_ Bohr effect (Kilmartin and Rossi-Bernard, 1971). Carbamylation of lysine residues has also been established in RuBisco (Lundqvist and Schneider, 1991), a key enzyme for photosynthetic carbon fixation, and in microbial beta-lactamases (Golemi et al., 2001; Maveyraud et al., 2000). CO_2_-dependent carbamylation as a general and important post-translational protein modification involved in physiological regulation was first proposed by George Lorimer (1983). It is clear from the known examples, and systematic application of mass spectrometric tools (Linthwaite et al., 2018), that only specific lysine residues in some proteins are able to be carbamylated. Our data now suggests that CO_2_-carbamylation plays an important physiological role in the control of breathing.

The direct and independent role of CO_2_-sensing in central chemoreception was first suggested by Shams (Shams, 1985). Our new data provides unequivocal evidence to demonstrate a molecular mechanism and role for direct CO_2_ sensing in the control of breathing. This molecular mechanism and pathway functions independently from any secondary changes in pH that could result from the altered balance between CO_2_ and HCO3-during hypercapnia. Our data further suggest that the role of Cx26 and direct CO_2_ sensing is restricted to an area of the ventral medullary surface in the caudal brain stem - ventral and medial to the lateral reticular nucleus. This area corresponds to the classically described caudal chemosensing area (Schlaefke et al., 1970; Trouth et al., 1973a, b) and in particular a subregion that has been more recently studied and termed the “caudal parapyramidal” area. The caudal parapyramidal area contains serotonergic neurons that are highly pH sensitive (Ribas-Salgueiro et al., 2003; Ribas-Salgueiro et al., 2005). Interestingly, our lentiviral vector consistently transduced glial cells at the very ventral surface of the medulla in this region (Figure 7). Our previous work has shown, with the aid of a knock-in reporter, that Cx26 is expressed in GFAP+ cells with cell bodies at the very surface of the parenchyma (Figure 10 of (Huckstepp et al., 2010b)). Our current work is consistent with this finding and shows that these superficial GFAP+ cells have: a cell body with a flattened edge that forms part of the very surface of the marginal glial layer; and long dorsally directed processes that extended both rostrally and medially (Figures 5a and 6). The cell body is ideally placed to detect changes in PCO_2_ in the CSF, but their processes projecting both rostrally and medially could contact neurons of the preBötzinger complex to directly alter inspiratory activity, or neurons of the raphé obscurus or pallidus, which detect changes in pH and are part of the chemosensory network (Bradley et al., 2002; Wang et al., 2002; Wang et al., 2001).

**Figure 7.**
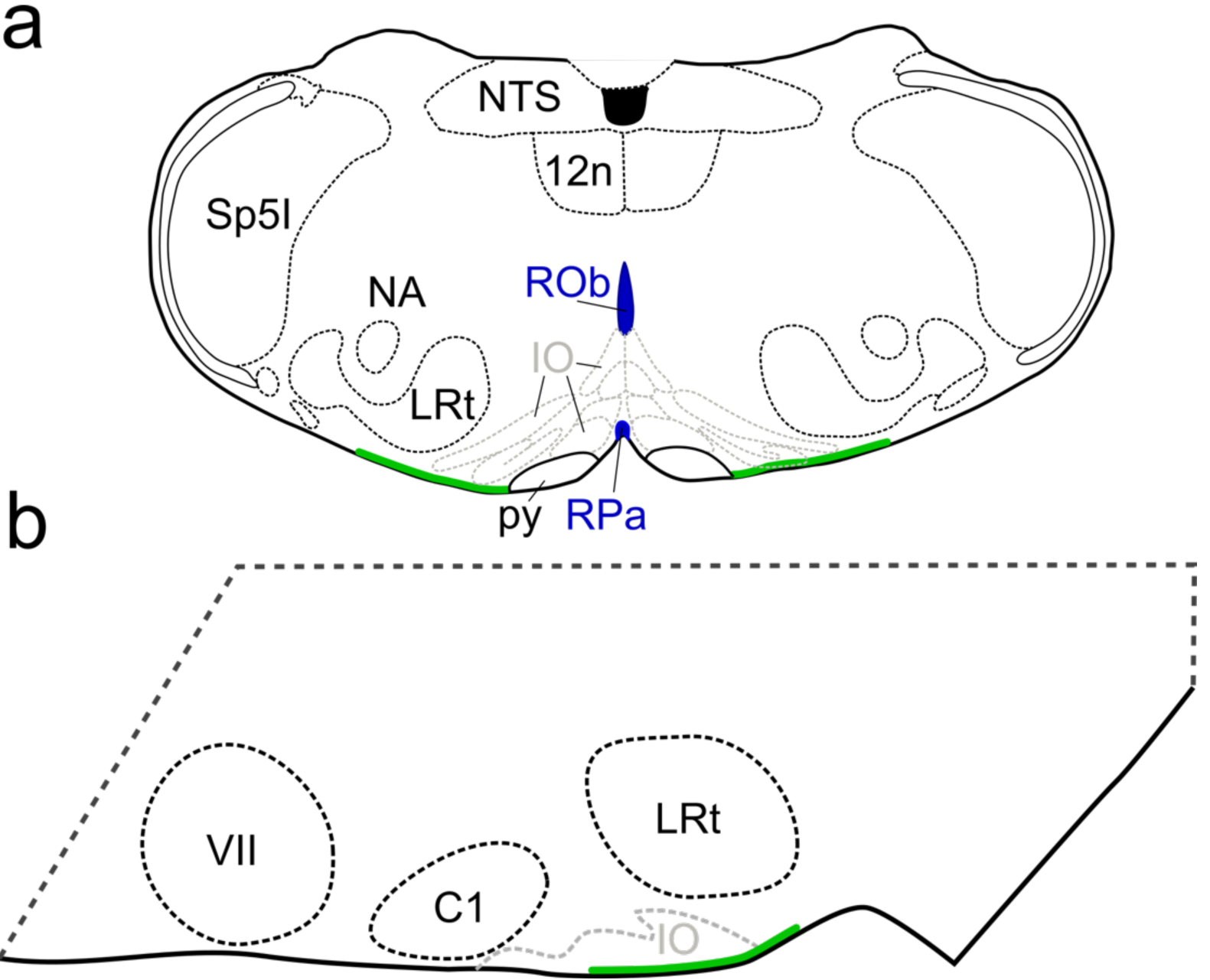
Schematic of the location of glial cells which, when transduced with Cx26^DN^ at the medullary surface, reduced the chemosensitivity of breathing. Location of cells is shown in the coronal (a) and parasagittal (b) planes. The cells are in an area ventral to the lateral reticular nucleus (LRt) and inferior olive (IO). The area reaches laterally to the same parasagittal plane as the nucleus ambiguus (NA) extends medially to the pyramids (py). NTS, nucleus tractus solitarius; 12n, hypoglossal nucleus; ROb, raphé obscurus; RPa, raphé pallidus; SP5I, spinal trigeminal nucleus interpolar part; VII, facial nucleus; C1, lateral paragigantocellular nucleus C1.

Our findings are robust and reproducible. We discovered the importance of Cx26 in the caudal parapyramidal area in a pilot experiment (Supplementary Figure 5), and used these data to design a properly powered experiment to test the importance of Cx26 in this area. For practical reasons, the groups of mice for the experiment reported in Figure 5 were divided into sub-cohorts of 6 per treatment group. In each of these sub-cohorts, the effect of Cx26^DN^ replicated that seen in the pilot experiment. We also observed the effect of Cx26^DN^ at 1.5, 2 and 2.5 weeks, however by 3 weeks post-transduction the effect of Cx26^DN^ on breathing had disappeared. This is presumably due to compensation within the respiratory network, which is known to be highly plastic (Johnson and Mitchell, 2013). For example: the effects of carotid body denervation are largely reversed over the course of 2 weeks (Pan et al., 1998); mice engineered to express a mutant Phox2b gene selectively in neurons of the RTN have no CO_2_ chemosensitivity for the first 9 days postnatally, but recover about 40% of their CO_2_ chemosensitivity by 4 months (Ramanantsoa et al., 2011). Interestingly, the recovered chemosensitivity in these mice was mediated almost exclusively via an adaptive increase in tidal volume, rather than frequency (Ramanantsoa et al., 2011) - it is therefore tempting to speculate that this could have arisen via the progressive postnatal expression of Cx26 in glial cells of the caudal parapyramidal area. There are several ways that the respiratory and chemosensory networks could compensate for the effects of Cx26^DN^, the most obvious being via strengthening of other chemosensory inputs such as those from the raphé, RTN or carotid bodies. However, a more subtle compensatory mechanism might be through upregulation of ATP receptors on the neurons downstream from the chemosensory glia in the caudal parapyramidal area. It is unlikely that we transduced every chemosensitive glial cell in this area, hence an upregulation of ATP receptors could maximise the effects of the non-transduced glial cells that remain chemosensitive.

Over-expression of Cx26^WT^ by itself did not enhance the chemosensitivity of breathing. Our construct drives expression in GFAP+ cells, which could either be part of the chemosensory network, or outside of it. For those glial cells within the chemosensory network it may be that the endogenous levels of expression are sufficient and additional expression of Cx26 can give no further gain of chemosensitivity - a “ceiling” effect. Cx26^WT^ expression in chemosensory cells outside of the network may be ineffective in enhancing chemosensitivity because these cells do not project to (and release ATP in) the correct locations to alter breathing.

### Gap junctions or hemichannels?

Cx26^DN^ and Cx26^WT^ colocalise to gap junction plaques and the FRET analysis shows that they coassemble into heteromeric gap junctions (Figure 2). By itself, Cx26^DN^ forms functional gap junctions which allow dye transfer (Supplementary Figure 10). Cx26^WT^ gap junctions can be closed by CO_2_ acting via the same carbamylation motif that opens the hemichannel (Nijjar et al., 2019) and Cx26^DN^ subunits form gap junctions that are insensitive to CO_2_ (Supplementary Figure 10). We therefore expect that Cx26^DN^ would have a similar dominant negative action on the CO_2_ sensitivity of endogenous Cx26 gap junctions and cannot exclude that Cx26^DN^-mediated loss of CO_2_-dependent gap junction closure could contribute to the observed results. However, there is abundant evidence for the presence of Cx26 hemichannels (Huckstepp et al., 2010b), and it is mechanistically simpler to understand how the CO_2_-gated release of ATP via Cx26 hemichannels could result in enhanced neuronal excitation and hence the adaptive changes in ventilation, compared to CO_2_-gated loss of gap junction communication between glial cells.

### Relationship with other chemosensory areas in the medulla oblongata

The RTN has received extensive attention as a chemosensory area (Burke et al., 2015; Guyenet et al., 2010; Kumar et al., 2015; Mulkey et al., 2004; Wang et al., 2013b). It is interesting that Cx26^DN^ expression in the RTN had no effect on chemosensitivity of breathing. CO_2_-dependent ATP release (most probably via Cx26) has been observed *in vivo* from the rostral ventral surface of the medulla, but this is more medial and probably closer to the Raphé magnus rather than the RTN, which is significantly more lateral (Gourine et al., 2005; Huckstepp et al., 2010b). Our data would suggest that chemosensory mechanisms within the RTN are completely independent of Cx26 and may not involve the direct sensing of CO_2_. The observation of pH dependent ATP release from RTN astrocytes (Gourine et al., 2010) via a membrane process involving the sodium bicarbonate transporter (NBCe1) and reversed Na^+^-Ca^2+^ exchange (Turovsky et al., 2016) supports this hypothesis.

There is compelling evidence to support the serotonergic medullary raphé neurons as an important mediator of respiratory chemosensitivity. Selective DREADD-mediating silencing of these neurons gave a substantial reduction (∼40%) of the adaptive change in ventilation to a 5% inspired CO_2_ challenge (Ray et al., 2011). Recently, chemosensory responses in RTN neurons have been shown to depend partially on serotonergic inputs from raphé neurons and RTN neurons have been proposed as relays of chemosensory information rather than primary chemosensors (Wu et al., 2019). Interestingly, the serotonergic raphé system extends caudally and will be medial to the chemosensory glial cells that we have identified in this study. As the surface glial cells that we transduced with Cx26^DN^ have long processes that project both rostrally and medially it is plausible that these cells could excite these raphé neurons via release of ATP or other gliotransmitters. Furthermore, there are additional pH-sensitive serotonergic neurons in the caudal parapyramidal area (Ribas-Salgueiro et al., 2003) that are likely to be intermingled with the glial cells that we describe in this study. CO_2_-dependent ATP release from these glial cells could potentially excite the neighbouring serotonergic neurons. As these serotonergic neurons project widely to other chemosensory areas of the medulla including the raphé obscurus, raphé pallidus, the NTS (Ribas-Salgueiro et al., 2005), their potential activation via release of ATP from CO_2_-sensitive glia in the same region gives a further mechanism by which this chemosensory signal could converge with that mediated via other chemosensory neurons (including the pH-dependent chemosensory pathway) and be propagated within the brain stem neural networks to facilitate breathing.

### Physiological significance of CO_2_ sensing versus other chemosensory mechanisms

Peripheral (mainly carotid body) and central chemoreceptors mediate the CO_2_-dependent control of breathing. Overall about 30% of the total chemosensory response is mediated peripherally and the remainder via central chemoreceptors (Heeringa et al., 1979). Additionally, there are two components to the adaptive changes in ventilation - an increase in respiratory frequency, and an increase in tidal volume (West, 2012). These two components combine to increase the rate of ventilation of the lungs: minute ventilation. Broadly, peripheral chemoreceptors appear primarily to increase respiratory frequency, whereas activation of central chemoreceptors have a more powerful effect on tidal volume (Li et al., 1999; Nattie and Li, 2001; Schlaefke et al., 1970). However, ATP release from astrocytes in the preBotzinger complex does modulate changes in respiratory frequency in response to elevated CO_2_ (Sheikhbahaei et al., 2018).

The widely accepted explanation for the chemosensitivity of breathing posits that pH sensitive neurons detect changes in the pH of arterial blood and then excite the medullary networks that control breathing. It is clear that the peripheral chemoreceptors are exclusively sensitive to pH (Schuitmaker et al., 1986). pH-sensitive TASK-1 channels contribute to the detection of pH changes in blood in the carotid body glomus cells and thus contribute to peripheral chemoreception (Trapp et al., 2008). Centrally, neurons from the RTN (Mulkey et al., 2004), the caudal parapyramidal area (Ribas-Salgueiro et al., 2003) and the medullary raphé (Richerson et al., 2001; Wang et al., 2001) respond to changes in pH most probably via TASK-2 (Gestreau et al., 2010; Koizumi et al., 2010; Kumar et al., 2015; Niemeyer et al., 2010; Pena-Munzenmayer et al., 2014; Wang et al., 2013a) and potentially GPR4 (Hosford et al., 2018; Kumar et al., 2015).

Given this multiplicity of evidence supporting the role of pH detection on regulation of breathing, why should CO_2_ sensing via Cx26 be particularly significant? Firstly, the contribution of central direct CO_2_ sensing to the regulation of breathing is about 50% of the centrally-generated chemosensory response (Shams, 1985). Viral transduction by Cx26^DN^ reduced the mean adaptive ventilatory response to 6% inspired CO_2_ by about 30% - mainly via a reduction of the increase in tidal volume. It is unlikely that we transduced all of the chemosensitive glial cells in this area, so this may be an underestimate of the contribution of this mechanism. Pharmacological blockade of Cx26 in the medulla, with imperfectly selective and potent agents, gave a reduction in the adaptive ventilatory response in anesthetized and ventilated rats of about 20% (Huckstepp et al., 2010b). The Cx26^DN^ transduction is clearly more effective than the prior pharmacological approach - it is highly selective and will completely remove Cx26-mediated CO_2_ sensitivity from the glial cells in which it is expressed.

As central chemoreceptors mediate about 70% of the adaptive response, CO_2_ sensing via Cx26 in the caudal parapyramidal area mediates just under half of the total central ventilatory response to modest levels of hypercapnia. A further comparison to give physiological context is that this contribution from Cx26 is slightly smaller than, but broadly comparable to, the DREADD-mediated inactivation of the entire population of raphé serotonergic neurons, which gave a 40% reduction in the adaptive ventilatory response at similar levels of inspired CO_2_ (Ray et al., 2011). The physiological importance of this contribution to chemosensing is further confirmed by the evolutionary conservation of the carbamylation motif and CO_2_-sensitivity in Cx26 in all amniotes for >400 million years (Dospinescu et al., 2019).

The contribution of Cx26 to respiratory chemosensitivity occurred over an intermediate (∼6%), but physiologically important, range of inspired CO_2_. There was no detectable effect of Cx26^DN^ on the ventilatory response to 3% inspired CO_2_. This is unlikely to reflect the properties of Cx26, as it is sensitive to changes in PCO_2_ from 20 to 70 mmHg (de Wolf et al., 2017; Huckstepp et al., 2010a), so would very likely be capable of detecting the small change in systemic PCO_2_ resulting from a 3% CO_2_ challenge. Cx26 contributes only to the adaptive change in tidal volume during hypercapnia (Figure 5). The ventilatory response to 3% CO_2_ involved only a small change in tidal volume (an increase of 0.34 µl.g^-1^ over V_T_ at 0% CO_2_ in Cx26^WT^ injected mice, Figure 5, Supplementary Table 1). If Cx26 contributed about 30% of this increase, the expected difference in the V_T_ between Cx26^WT^- and Cx26^DN^-injected mice would be ∼0.1 µl.g^-1^. Although there is a trend towards a reduced change in V_T_ at 3% CO_2_ (Figure 5, Supplementary Table 1), our experiments were insufficiently powered to detect such a small difference and hence cannot reveal any possible contribution of Cx26 at this level of inspired CO_2_. A further factor in determining the overall contribution of cells that express Cx26 to respiratory control, will be the way they and other populations of chemosensory cells, both central and peripheral, connect to the neuronal circuits controlling breathing.

At 9% inspired CO_2_, Cx26-mediated chemosensing makes no contribution to the regulation of breathing, and the ventilatory responses to these higher levels are most likely exclusively mediated by changes in pH. This may be because the acidification caused by the higher level of inspired CO_2_ can counteract the CO_2_-dependent opening of Cx26 hemichannels. Cx26 hemichannels are closed by strong acidification (Yu et al., 2007). We have previously reported that acidification reduces the conductance change evoked by a PCO_2_ stimulus in Cx26-expressing HeLa cells (Huckstepp et al., 2010a). Furthermore, the same increase in PCO_2_ at pH 7.35 evokes about half the ATP release from the chemosensory cells in the ventral medulla compared to that evoked at pH 7.5 (Huckstepp et al., 2010b). This is presumably because acidification makes carbamylation of the critical lysine residue harder to achieve (Meigh et al., 2013).

The caudal parapyramidal area (Figure 7) was the only location in which we found a contribution of Cx26 contributed to respiratory chemosensitivity. This area has been previously described as containing pH sensitive serotonergic neurons (Ribas-Salgueiro et al., 2003). The caudal parapyramidal area is thus sensitive to both chemosensory stimuli and is a potential point of convergence of the CO_2_-mediated and pH-mediated chemosensory signals. In the rostral medulla, our injections of Cx26^DN^ were too lateral to test whether Cx26 expression in more medial glial cells might also contribute to activation of the pH-sensitive raphé magnus neurons, but this would be an interesting hypothesis to investigate.

To conclude, the development of Cx26^DN^ as a tool to remove CO_2_ sensitivity from endogenously expressed Cx26 has provided the first genetic evidence for the involvement of Cx26 in the control of breathing. Our data provides a mechanistic link between the binding of CO_2_ to a structural motif on Cx26 and shows that direct sensing of CO_2_ by glial cells in a circumscribed area of the ventral medulla contribute nearly half of the centrally generated adaptive ventilatory response to CO_2_.

## Methods

### Animals

All animal procedures were evaluated by the Animal Welfare and Ethical Review Board of the University of Warwick and carried out in strict accordance with the Animals (1986) Scientific Procedures Act of the UK under the authority of Licence PC07DE9A3. Mice were randomly assigned to their groups (Cx26^WT^, Cx26^DN^ and sham). A total of 80 mice were used in this study.

#### Pilot experiment, and RTN

Male and female mice 12-20 weeks old were used. The mice had a floxed Cx26 allele on a C57BL6 background (EMMA strain 00245). These mice were not crossed with any cre lines so had normal wild type expression of Cx26. *Main caudal parapyramidal experiment*: C57BL6 WT Male mice 12-17 weeks were used. *Very caudal experiment*: C57BL6 WT Male mice 12-20 weeks were used.

### Cell lines

HeLa DH (obtained from ECACC) and stable Cx26-expressing HeLa cells (gift from K. Willecke) were maintained in Dulbecco’s modified Eagle’s medium (DMEM) supplemented with 10% fetal calf serum (FCS), 50 μg/mL Penicillin Streptomycin, and 3mM CaCl2 at 37°C.

### Construction of connexin gene constructs

Connexin43 DNA sequence from *Rattus norvegicus* was purchased from Addgene (mCherry-Cx43-7, plasmid #55023, gifted by Michael Davidson) and subcloned into a PUC 19 vector such that the transcript would form a fusion protein with whichever fluorophore we engineered to be 3’ (downstream) of Cx43 (as with Cx26 constructs).

Dominant Negative mutant Cx26 (Cx26^DN^) DNA - with R104A K125R mutations was produced through two steps. First, Cx26 DNA with the K125R mutation and omitted STOP codon (to allow for fusion proteins) was synthesised by Genscript USA from the Cx26 sequence (accession number NM_001004099.1), and subsequently subcloned into a PUC 19 vector such that the transcript would form a fusion protein with mCherry at the C-terminus of Cx26. Second, the R104A mutation was introduced by Agilent Quikchange site directed mutagenesis, using PUC19-Cx26(K125R) DNA as the template. Primer sequences for the mutagenesis were as follows: Cx26(R104A) forward 5’GGC CTA CCG GAG ACA CGA AAA GAA AGC GAA GTT CAT GAA GG 3’, Cx26(R104A) reverse 5’CCT TCA TGA ACT TCG CTT TCT TTT CGT GTC TCC GGT AGG CC 3’.

Design and characterization of the Clover and mRuby2 protein fluorophores used in these experiments was published by (Lam et al., 2012). mRuby2 DNA was purchased from Addgene (mRuby2-C1, plasmid #54768, gifted by Michael Davidson) and subcloned into a PUC 19 vector so that it was 3’ of whichever connexin we engineered to be 5’ (upstream) of mRuby2. Clover DNA was a gift from Sergey Kasparov, Bristol, and was subcloned into a PUC 19 vector so that it was 3’ of whichever connexin we engineered to be 5’ (upstream) of Clover.

### HeLa cell transfection

To express the desired constructs, HeLa cells were transiently transfected with 0.5μg of DNA of each pCAG-Connexin-fluorophore construct to be co-expressed (1μg total), using the GeneJuice transfection agent protocol (Merck Millipore).

### Experimental aCSF solutions

#### Control (35 mmHg PCO_2_)

124 mM NaCl, 26 mM NaHCO_3_, 1.25 mM NaH_2_PO_4_, 3 mM KCl, 10 mM D-glucose, 1 mM MgSO_4_, 2 mM CaCl_2_. This was bubbled with 95%O_2_ / 5% CO_2_ and had a final pH of ∼7.4.

#### Hypercapnic (55 mmHg PCO_2_)

100 mM NaCl, 50 mM NaHCO_3_, 1.25 mM NaH_2_PO_4_, 3 mM KCl, 10 mM D-glucose, 1 mM MgSO_4_, 2 mM CaCl_2_. This was bubbled with sufficient CO_2_ (approximately 9%, balance O_2_) to give a final pH of ∼7.4.

#### Zero Ca^2+^

124 mM NaCl, 26 mM NaHCO_3_, 1.25 mM NaH_2_PO_4_, 3 mM KCl, 10 mM D-glucose, 1 mM MgSO_4_, 2 mM MgCl_2_, 1mM EGTA. This was bubbled with 95%O_2_ / 5% CO_2_ and had a final pH of ∼7.4.

### Dye loading assay

We used a dye loading protocol that has been developed and extensively described in our prior work (Dospinescu et al., 2019; Huckstepp et al., 2010a; Meigh et al., 2015; Meigh et al., 2013; Meigh et al., 2014). HeLa-DH cells were plated onto coverslips and transfected with Cx26^DN^. Dye loading experiments were performed 24-72h after transfection. For experiments involving co-expression of Cx26^DN^ and Cx26^WT^, HeLa cells that stably expressed Cx26^WT^ were transfected with Cx26^DN^. Dye loading was performed over a 3 day period, beginning at 4 days post-transfection.

To perform the dye loading, cells were washed in control aCSF were then exposed to either control or hypercapnic solution containing 200 μM 5(6)-carboxyfluorescein (CBF) for 10 min. Subsequently, cells were returned to control solution with 200 μM CBF for 5 min, before being washed in control solution without CBF for 30-40 min to remove excess extracellular dye. A replacement coverslip of HeLa cells was used for each condition. For each coverslip, mCherry staining was imaged to verify Cx26 expression.

To quantify fluorescence intensity, cells were imaged by epifluorescence (Scientifica Slice Scope (Scientifica Ltd, Uckfield, UK), Cairn Research OptoLED illumination (Cairn Research Limited, Faversham, UK), 60x water Olympus immersion objective, NA 1.0 (Scientifica), Hamamatsu ImageEM EMCCD camera (Hamamatsu Photonics K.K., Japan), Metafluor software (Cairn Research). Using ImageJ, regions of interest (ROI) were drawn around individual cells and the mean pixel intensity for each ROI obtained. The mean pixel intensity of the background fluorescence was also measured in a representative ROI, and this value was subtracted from the measures obtained from the cells. This procedure was used to subtract the background fluorescence from every pixel of all of the images displayed in the figures. At least 40 cells were measured in each condition, and the mean pixel intensities plotted as cumulative probability distributions.

The experiments were replicated independently (independent transfections) at least 5 times for each Cx26 variant and condition. All experiments performed at room temperature.

### FRET data capture and analysis

For FRET analysis, HeLa cells co-transfected with Cx-Clover (donor) and Cx-mRuby2 (acceptor) 72 hours after transfection were washed 3x with PBS, fixed with 4% PFA (paraformaldehyde) for 20-30 minutes, washed a further 3x with PBS, and then stored in PBS at 4^0^C. FRET studies were carried out within 2 weeks of fixation. They examined with a Zeiss LSM 710 Confocal microscope; C-Apochromat 63x/1.20 W Korr M27. Two channels were recorded: 495-545nm (clover) and 650-700nm (mRuby2), and images were acquired sequentially with 458nm and 561nm argon lasers (respectively). Imaging parameters for the Clover channel are as follows: power, 30.0; pinhole, 78.5; gain (master), 700; digital offset, 0; digital gain, 1.0. Imaging parameters for the mRuby2 channel are as follows: power, 30.0; pinhole, 78.5; gain (master), 750; digital offset, 0; digital gain, 1.0. Photobleaching was performed using the 561nm laser (as it only excites mRuby2) for 80 frames at 100% power, targeting ROI’s. Image acquisition was as follows: ROIs were selected and drawn, including a background region and an ROI that was not to be bleached; pre-bleaching images were acquired for each channel; mRuby2 was photobleached; post-bleaching images were acquired for each channel. Acquisition parameters were kept identical across samples to allow comparison of results.

Pixel intensities from each background-adjusted ROI were used to calculate FRET efficiency (E), bleaching efficiency (B), relative donor abundance (D), relative acceptor abundance (A), relative donor-acceptor ratio (DA ratio), relative acceptor quantity (A level), and donor-acceptor distance (R) as follows:

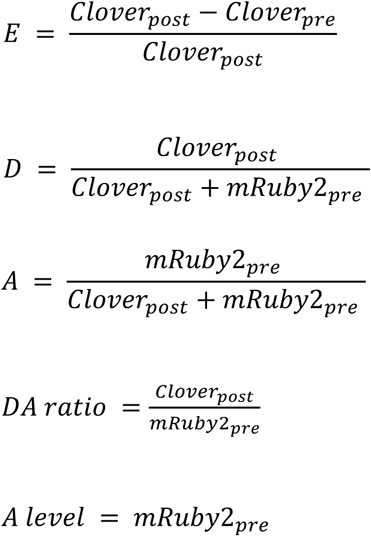

Colocalisation threshold calculations were carried out in ImageJ using the Costes method (Costes et al., 2004); 100 iterations, omitting zero-zero pixels in threshold calculation.

### Lentivirus design and production

Two lentivirus (LV) constructs were designed to introduce either the Cx26DN or Cx26WT gene into the host cell genome (Supplementary Figure 3). Constructs are ∼4900bp in length. The sequence of interest consisted off the Cx26 gene variant (Cx26^WT^ or Cx26^DN^) immediately followed by an internal ribosome entry site and Clover sequence (Cx26:IRES:Clover). The IRES sequence was from the encephalomyocarditis virus (Bochkov and Palmenberg, 2006). Lentivirus constructs were produced and packaged by Cyagen Biosciences (USA) using the third-generation packing system. Constructs had a titre > 10^8^ TU/mL; as confirmed by qPCR on genomic DNA extracted from infected cells.

### Stereotaxic viral transduction

Anaesthesia was induced by inhalation of isoflurane (4%). The mouse was then placed on a thermocoupled heating pad (TCAT-2LV, Physitemp) to maintain body temperature at 33°C and headfixed into a stereotaxic frame. A face mask was used to maintain anesthesia (isoflurane, intranasal, 0.5-2.5%). Atropine was provided (sub-cutaneous, 0.05 mg/kg) before surgery to stop pleural effusion. Adequacy of anesthesia was assessed by respiratory rate, body temperature, and pedal withdrawal reflex. Pre-operative Meloxicam (sub-cutaneous, 2 mg/kg) and post-operative Buprenorphine (intraperitoneal, 0.05 mg/kg) were provided for analgesia. If any animal showed signs of pain in the days following surgery additional analgesia was administered as required. No animals displaying signs of pain or receiving analgesia were used in plethysmography recordings.

To maintain consistent placement of the injection pipette, the intra-aural line was adjusted so that bregma was level to a point on the skull 2 mm caudal to bregma. Two small holes were made in the interparietal plate to allow for bilateral injection of virus particles via a micropipette lowered into the correct position via a stereotaxic manipulator. The injections were performed manually, using a 1 ml syringe, at a rate ∼200 nl/min. A total of 350-400 nl of undiluted virus or saline was injected per side of the brain, the experimenter was blind to the injection solution. Co-ordinates (mm) relative to bregma were: *RTN:* 5.7 and 5.9 caudal (two injections per side of brain), 1.1 lateral and 5.3-5.9 ventral, injection arm at 9° to vertical; *caudal parapyramidal area*: 5.95 caudal, 0.8 lateral, and 5.7-6.1 ventral, injection arm at 0° to vertical; *very caudal area:* 6.2 caudal, 0.8 lateral and 5.7-6.2 ventral, injection arm at 0° to vertical; *pilot experiment*: 5.9 caudal, 1.1 lateral, and 5.5-6.0 ventral, injection arm at 0° to vertical. Co-ordinates were confirmed by *post hoc* immunostaining for viral driven expression of fluorophores. Data were only included from mice whose injection sites were within the correct location

### Immunohistochemistry

Mice were culled by overdose of Isoflurane or intraperitoneal injection of Sodium Pentobarbital (>100 mg/kg), and transcardially perfused with paraformaldehyde (PFA; 4%). The brain was harvested and post-fixed in 4% PFA for 24hrs at 4°C (to further increase tissue fixing), before being transferred to 30% sucrose (for cryoprotection) until the brain sunk to the bottom of the sucrose – usually ∼2-3 days. PFA and sucrose were made in PBS.

For sectioning, brainstems were isolated, mounted with Tissue-Tek optimum cutting temperature compound (Sakura Finetek), and cut either sagittally or coronally at 40 μm on a cryostat (Bright Instruments). Sections were arranged in order, in 24-well plates, and stored in PBS. For immunostaining, free-floating sections were incubated at room temperature overnight in PBS containing 0.1% Triton X-100 (Sigma-Aldrich, UK) (0.1% PBS-T) and the appropriate primary antibodies from: goat anti-Choline Acetyltransferase (ChAT) (1:100) (Merck UK, ab144), chicken anti-GFAP (glial fibrillary acidic protein) (1:500) (abcam, ab4674), rabbit anti-GFAP (1:500) (abcam, ab68428). Sections were then washed in PBS for 6 × 5 mins before incubation at room temperature for 2.5-4 hrs in 0.1% PBS-T containing the appropriate secondary antibodies from: Donkey anti-goat Alexa Fluor 594 (1:250) (abcam, ab150132), goat anti-rabbit Alexa Fluor 594 (1:250) (abcam, ab150080), goat anti-chicken Alexa Fluor 488 (1:250) (abcam, ab150169). After secondary antibody incubation, sections were again washed in PBS for 6 × 5 mins before mounting onto polylysine-coated slides (Polysine, VWR). Mounted slices were left to dehydrate overnight before applying coverslips using either Aqua-Poly/Mount (Polysciences Inc., Germany) or Fluoroshield with DAPI (Sigma-Aldrich, UK). All steps were performed at 21°C and during any period of incubation or washing sections were gently agitated on a shaker.

### Whole-body plethysmography

A custom built plethysmograph, constructed from a plexiglass box (0.5 l), was equiped with an air-tight lid, a pressure transducer to detect the breathing movements of the mouse, and gas inlets and outlets to permit gas flow through the chamber. A heated (via a water bath) mixture of O_2_ (∼20%), N_2_ (80%) and CO_2_ (0-9%) flowed through the chamber regulated to a rate of 1 l.min^-1^. The amount of O_2_ and CO_2_ in the mixture was measured, just prior to entry into the chamber, by a gas analyzer (Hitech Intruments, GIR250 Dual Sensor Gas Analyzer). Pressure signals were recorded with an NL108T2 – Disposable Physiological Pressure Transducer (Digitimer) amplified and filtered using the NeuroLog system connected to a 1401 interface and acquired on a computer using Spike2 software (Cambridge Electronic Design). Airflow measurements were used to calculate: tidal volume (V_T_: signal trough at the end of expiration subtracted from the peak signal during inspiration, converted to ml following calibration with a 1 ml syringe), respiratory frequency (f_R_: breaths per minute), and minute ventilation (V_E_) (calculated as V_T_ × f_R_). The temperature inside the plethysmograph was maintained at ∼31°C, thermoneutral for C57BL/6 mice. The experimenter was only unblinded to the identity of the injected virus once acquisition and analysis of all plethysomographic recordings had been performed.

## Statistics and reproducibility

For the dye-loading experiments, box-whisker plots were made and statistical tests performed on the median change in pixel intensity where each replicate was a different transfection of the cells. The statistical package R was used for this analysis. For the FRET data, linear regression analysis and plots were carried out in GraphPad Prism. The plethsymographic data was analysed in SPSS via a 2-way mixed effects, repeated measures ANOVA. Multiple post-hoc comparisons were checked by the false discovery rate method, with the maximum allowed false discovery rate set to 0.05 (Curran-Everett, 2000). All reported comparisons pass this test. Power calculations for the experiment in Figure 5 were based on the pilot data of Supplementary Figure 5, and were performed in GPower 3.1 for a 2-way ANOVA with repeated measures, within and between factor interactions. Further statistical details can be found in the relevant figure legends.

## Acknowledgements

We thank the MRC (MR/N003918/1) for support. JdW was a doctoral student supported by the MRC (MR/J003964/1). ND was a Royal Society Wolfson Research Merit Award Holder.

## Author contributions

Molecular biology: JdW, LM, JC

*In vitro* experiments: LM, JdW, SN

Immunocytochemical staining: JdW, AB

Surgical procedures: JdW, AB, RTH

Plethysmography: JdW

Data analysis: JdW, LM, ND

Study design: ND, JdW, RTH

Writing paper: ND, JdW, RTH and AB

All authors commented on drafts of the paper.

## Data availability

All data generated or analysed during this study are included in this published article (and its supplementary information files).

## References

Bernard, D.G., Li, A., and Nattie, E.E. (1996). Evidence for central chemoreception in the midline raphe. J Appl Physiol 80, 108–115.

Bochkov, Y.A., and Palmenberg, A.C. (2006). Translational efficiency of EMCV IRES in bicistronic vectors is dependent upon IRES sequence and gene location. Biotechniques 41, 283-284, 286, 288 passim.

Bradley, S.R., Pieribone, V.A., Wang, W., Severson, C.A., Jacobs, R.A., and Richerson, G.B. (2002). Chemosensitive serotonergic neurons are closely associated with large medullary arteries. Nat Neurosci 5, 401–402.

Brust, R.D., Corcoran, A.E., Richerson, G.B., Nattie, E., and Dymecki, S.M. (2014). Functional and developmental identification of a molecular subtype of brain serotonergic neuron specialized to regulate breathing dynamics. Cell Rep 9, 2152–2165.

Burke, P.G., Kanbar, R., Viar, K.E., Stornetta, R.L., and Guyenet, P.G. (2015). Selective optogenetic stimulation of the retrotrapezoid nucleus in sleeping rats activates breathing without changing blood pressure or causing arousal or sighs. J Appl Physiol (1985) 118, 1491–1501.

Costes, S.V., Daelemans, D., Cho, E.H., Dobbin, Z., Pavlakis, G., and Lockett, S. (2004). Automatic and quantitative measurement of protein-protein colocalization in live cells. Biophys J 86, 3993–4003.

Curran-Everett, D. (2000). Multiple comparisons: philosophies and illustrations. Am J Physiol Regul Integr Comp Physiol 279, R1–8.

de Wolf, E., Cook, J., and Dale, N. (2017). Evolutionary adaptation of the sensitivity of connexin26 hemichannels to CO2. Proc R Soc B 284: 20162723.

Di, W.L., Gu, Y., Common, J.E., Aasen, T., O’Toole, E.A., Kelsell, D.P., and Zicha, D. (2005). Connexin interaction patterns in keratinocytes revealed morphologically and by FRET analysis. J Cell Sci 118, 1505–1514.

Dospinescu, V.-M., Nijjar, S., Spanos, F., Cook, J., de Wolf, E., Biscotti, M.A., Gerdol, M., and Dale, N. (2019). Structural determinants of CO2-sensitivity in the β connexin family suggested by evolutionary analysis. Communications Biology 2, 331.

Eldridge, F.L., Kiley, J.P., and Millhorn, D.E. (1985). Respiratory responses to medullary hydrogen ion changes in cats: different effects of respiratory and metabolic acidoses. J Physiol 358, 285–297.

Forster, H.V. (2018). Julius H. Comroe Distinguished Lecture: Interdependence of neuromodulators in the control of breathing. J Appl Physiol (1985) 125, 1511–1525.

Gestreau, C., Heitzmann, D., Thomas, J., Dubreuil, V., Bandulik, S., Reichold, M., Bendahhou, S., Pierson, P., Sterner, C., Peyronnet-Roux, J., et al. (2010). Task2 potassium channels set central respiratory CO2 and O2 sensitivity. Proc Natl Acad Sci U S A 107, 2325–2330.

Golemi, D., Maveyraud, L., Vakulenko, S., Samama, J.P., and Mobashery, S. (2001). Critical involvement of a carbamylated lysine in catalytic function of class D beta-lactamases. Proc Natl Acad Sci U S A 98, 14280–14285.

Gourine, A.V., Kasymov, V., Marina, N., Tang, F., Figueiredo, M.F., Lane, S., Teschemacher, A.G., Spyer, K.M., Deisseroth, K., and Kasparov, S. (2010). Astrocytes control breathing through pH-dependent release of ATP. Science 329, 571–575.

Gourine, A.V., Llaudet, E., Dale, N., and Spyer, K.M. (2005). ATP is a mediator of chemosensory transduction in the central nervous system. Nature 436, 108–111.

Guyenet, P.G. (2008). The 2008 Carl Ludwig Lecture: retrotrapezoid nucleus, CO2 homeostasis, and breathing automaticity. J Appl Physiol 105, 404–416.

Guyenet, P.G., Bayliss, D.A., Mulkey, D.K., Stornetta, R.L., Moreira, T.S., and Takakura, A.T. (2008). The retrotrapezoid nucleus and central chemoreception. Adv Exp Med Biol 605, 327–332.

Guyenet, P.G., Stornetta, R.L., and Bayliss, D.A. (2010). Central respiratory chemoreception. The Journal of comparative neurology 518, 3883–3906.

Heeringa, J., Berkenbosch, A., de Goede, J., and Olievier, C.N. (1979). Relative contribution of central and peripheral chemoreceptors to the ventilatory response to CO2 during hyperoxia. Respir Physiol 37, 365–379.

Hosford, P.S., Mosienko, V., Kishi, K., Jurisic, G., Seuwen, K., Kinzel, B., Ludwig, M.G., Wells, J.A., Christie, I.N., Koolen, L., et al. (2018). CNS distribution, signalling properties and central effects of G-protein coupled receptor 4. Neuropharmacology 138, 381–392.

Huckstepp, R.T., and Dale, N. (2011). Redefining the components of central CO2 chemosensitivity--towards a better understanding of mechanism. J Physiol 589, 5561–5579.

Huckstepp, R.T., Eason, R., Sachdev, A., and Dale, N. (2010a). CO2-dependent opening of connexin 26 and related beta connexins. J Physiol 588, 3921–3931.

Huckstepp, R.T., id Bihi, R., Eason, R., Spyer, K.M., Dicke, N., Willecke, K., Marina, N., Gourine, A.V., and Dale, N. (2010b). Connexin hemichannel-mediated CO2-dependent release of ATP in the medulla oblongata contributes to central respiratory chemosensitivity. J Physiol 588, 3901–3920.

Johnson, R.A., and Mitchell, G.S. (2013). Common mechanisms of compensatory respiratory plasticity in spinal neurological disorders. Respir Physiol Neurobiol 189, 419–428.

Kilmartin, J.V., and Rossi-Bernard, L. (1971). The binding of carbon dioxide by horse haemoglobin. Biochemical journal 124, 31–45.

Koizumi, H., Smerin, S.E., Yamanishi, T., Moorjani, B.R., Zhang, R., and Smith, J.C. (2010). TASK Channels Contribute to the K+-Dominated Leak Current Regulating Respiratory Rhythm Generation In Vitro. J. Neurosci. 30, 4273–4284.

Kumar, N.N., Velic, A., Soliz, J., Shi, Y., Li, K., Wang, S., Weaver, J.L., Sen, J., Abbott, S.B., Lazarenko, R.M., et al. (2015). Regulation of breathing by CO(2) requires the proton-activated receptor GPR4 in retrotrapezoid nucleus neurons. Science 348, 1255–1260.

Lam, A.J., St-Pierre, F., Gong, Y., Marshall, J.D., Cranfill, P.J., Baird, M.A., McKeown, M.R., Wiedenmann, J., Davidson, M.W., Schnitzer, M.J., et al. (2012). Improving FRET dynamic range with bright green and red fluorescent proteins. Nat Methods 9, 1005–1012.

Li, A., Randall, M., and Nattie, E.E. (1999). CO(2) microdialysis in retrotrapezoid nucleus of the rat increases breathing in wakefulness but not in sleep. J Appl Physiol 87, 910–919.

Linthwaite, V.L., Janus, J.M., Brown, A.P., Wong-Pascua, D., O’Donoghue, A.C., Porter, A., Treumann, A., Hodgson, D.R.W., and Cann, M.J. (2018). The identification of carbon dioxide mediated protein post-translational modifications. Nat Commun 9, 3092.

Liu, B., Paton, J.F., and Kasparov, S. (2008). Viral vectors based on bidirectional cell-specific mammalian promoters and transcriptional amplification strategy for use in vitro and in vivo. BMC Biotechnol 8, 49.

Loeschcke, H.H. (1982). Central chemosensitivity and the reaction theory. J Physiol 332, 1–24.

Lorimer, G.H. (1983). Carbon-Dioxide and Carbamate Formation - the Makings of a Biochemical Control-System. Trends in Biochemical Sciences 8, 65–68.

Lundqvist, T., and Schneider, G. (1991). Crystal structure of the ternary complex of ribulose-1,5-bisphosphate carboxylase, Mg(II), and activator CO2 at 2.3-A resolution. Biochemistry 30, 904–908.

Maveyraud, L., Golemi, D., Kotra, L.P., Tranier, S., Vakulenko, S., Mobashery, S., and Samama, J.P. (2000). Insights into class D beta-lactamases are revealed by the crystal structure of the OXA10 enzyme from Pseudomonas aeruginosa. Structure 8, 1289–1298.

Meigh, L., Cook, D., Zhang, J., and Dale, N. (2015). Rational design of new NO and redox sensitivity into connexin26 hemichannels. Open Biol 5, 140208.

Meigh, L., Greenhalgh, S.A., Rodgers, T.L., Cann, M.J., Roper, D.I., and Dale, N. (2013). CO2 directly modulates connexin 26 by formation of carbamate bridges between subunits. eLife 2, e01213.

Meigh, L., Hussain, N., Mulkey, D.K., and Dale, N. (2014). Connexin26 hemichannels with a mutation that causes KID syndrome in humans lack sensitivity to CO2. eLife 3, e04249.

Mitchell, R.A., Loeschcke, H.H., Severinghaus, J.W., Richardson, B.W., and Massion, W.H. (1963). Regions of respiratory chemosensitivity on the surface of the medulla. Ann N Y Acad Sci 109, 661–681.

Mulkey, D.K., Mistry, A.M., Guyenet, P.G., and Bayliss, D.A. (2006). Purinergic P2 receptors modulate excitability but do not mediate pH sensitivity of RTN respiratory chemoreceptors. J Neurosci 26, 7230–7233.

Mulkey, D.K., Stornetta, R.L., Weston, M.C., Simmons, J.R., Parker, A., Bayliss, D.A., and Guyenet, P.G. (2004). Respiratory control by ventral surface chemoreceptor neurons in rats. Nat Neurosci 7, 1360–1369.

Muller, D.J., Hand, G.M., Engel, A., and Sosinsky, G.E. (2002). Conformational changes in surface structures of isolated connexin 26 gap junctions. EMBO J 21, 3598–3607.

Nattie, E.E., Fung, M.L., Li, A., and St John, W.M. (1993). Responses of respiratory modulated and tonic units in the retrotrapezoid nucleus to CO2. Respir Physiol 94, 35–50.

Nattie, E.E., and Li, A. (2001). CO2 dialysis in the medullary raphe of the rat increases ventilation in sleep. J Appl Physiol 90, 1247–1257.

Niemeyer, M.I., Cid, L.P., Pena-Munzenmayer, G., and Sepulveda, F.V. (2010). Separate gating mechanisms mediate the regulation of K2P potassium channel TASK-2 by intra- and extracellular pH. J Biol Chem 285, 16467–16475.

Nijjar, S., Maddison, D., Meigh, L., de Wolf, E., Rodgers, T., Cann, M., and Dale, N. (2019). Opposing modulation of Cx26 gap junctions and hemichannels by CO2. bioRxiv, 584722.

Pan, L.G., Forster, H.V., Martino, P., Strecker, P.J., Beales, J., Serra, A., Lowry, T.F., Forster, M.M., and Forster, A.L. (1998). Important role of carotid afferents in control of breathing. J Appl Physiol (1985) 85, 1299–1306.

Pena-Munzenmayer, G., Niemeyer, M.I., Sepulveda, F.V., and Cid, L.P. (2014). Zebrafish and mouse TASK-2 K(+) channels are inhibited by increased CO2 and intracellular acidification. Pflugers Arch 466, 1317–1327.

Ramanantsoa, N., Hirsch, M.-R., Thoby-Brisson, M., Dubreuil, V., Bouvier, J., Ruffault, P.-L., Matrot, B., Fortin, G., Brunet, J.-F., Gallego, J., et al. (2011). Breathing without CO2 Chemosensitivity in Conditional Phox2b Mutants. The Journal of Neuroscience 31, 12880–12888.

Ray, R.S., Corcoran, A.E., Brust, R.D., Kim, J.C., Richerson, G.B., Nattie, E., and Dymecki, S.M. (2011). Impaired respiratory and body temperature control upon acute serotonergic neuron inhibition. Science 333, 637–642.

Ribas-Salgueiro, J.L., Gaytan, S.P., Crego, R., Pasaro, R., and Ribas, J. (2003). Highly H+-sensitive neurons in the caudal ventrolateral medulla of the rat. J Physiol 549, 181–194.

Ribas-Salgueiro, J.L., Gaytán, S.P., Ribas, J., and Pásaro, R. (2005). Characterization of efferent projections of chemosensitive neurons in the caudal parapyramidal area of the rat brain. Brain research bulletin 66, 235–248.

Richerson, G.B. (2004). Serotonergic neurons as carbon dioxide sensors that maintain pH homeostasis. Nat Rev Neurosci 5, 449–461.

Richerson, G.B., Wang, W., Tiwari, J., and Bradley, S.R. (2001). Chemosensitivity of serotonergic neurons in the rostral ventral medulla. Respir Physiol 129, 175–189.

Schlaefke, M.E., See, W.R., Herker-See, A., and Loeschcke, H.H. (1979). Respiratory response to hypoxia and hypercapnia after elimination of central chemosensitivity. Pflugers Arch 381, 241–248.

Schlaefke, M.E., See, W.R., and Loeschcke, H.H. (1970). Ventilatory response to alterations of H+ ion concentration in small areas of the ventral medullary surface. Respir Physiol 10, 198–212.

Schuitmaker, J.J., Berkenbosch, A., De Goede, J., and Olievier, C.N. (1986). Effects of CO2 and H+ on the ventilatory response to peripheral chemoreceptor stimulation. Respir Physiol 64, 69–79.

Shams, H. (1985). Differential effects of CO2 and H+ as central stimuli of respiration in the cat. J Appl Physiol 58, 357–364.

Sheikhbahaei, S., Turovsky, E.A., Hosford, P.S., Hadjihambi, A., Theparambil, S.M., Liu, B., Marina, N., Teschemacher, A.G., Kasparov, S., Smith, J.C., et al. (2018). Astrocytes modulate brainstem respiratory rhythm-generating circuits and determine exercise capacity. Nat Commun 9, 370.

Solomon, I.C., Halat, T.J., El-Maghrabi, M.R., and O’Neal, M.H., 3rd (2001). Localization of connexin26 and connexin32 in putative CO(2)-chemosensitive brainstem regions in rat. Respir Physiol 129, 101–121.

Trapp, S., Aller, M.I., Wisden, W., and Gourine, A.V. (2008). A role for TASK-1 (KCNK3) channels in the chemosensory control of breathing. J Neurosci 28, 8844–8850.

Trouth, C.O., Loeschcke, H.H., and Berndt, J. (1973a). Histological structures in the chemosensitive regions on the ventral surface of the cat’s medulla oblongata. Pflugers Arch 339, 171–183.

Trouth, C.O., Loeschcke, H.H., and Berndt, J. (1973b). A superficial substrate on the ventral surface of the medulla oblongata influencing respiration. Pflugers Arch 339, 135–152.

Turovsky, E., Theparambil, S.M., Kasymov, V., Deitmer, J.W., Del Arroyo, A.G., Ackland, G.L., Corneveaux, J.J., Allen, A.N., Huentelman, M.J., Kasparov, S., et al. (2016). Mechanisms of CO2/H+ Sensitivity of Astrocytes. J Neurosci 36, 10750–10758.

Valdez Capuccino, J.M., Chatterjee, P., Garcia, I.E., Botello-Smith, W.M., Zhang, H., Harris, A.L., Luo, Y., and Contreras, J.E. (2019). The connexin26 human mutation N14K disrupts cytosolic intersubunit interactions and promotes channel opening. J Gen Physiol 151, 328–341.

Wang, S., Benamer, N., Zanella, S., Kumar, N.N., Shi, Y., Bevengut, M., Penton, D., Guyenet, P.G., Lesage, F., Gestreau, C., et al. (2013a). TASK-2 channels contribute to pH sensitivity of retrotrapezoid nucleus chemoreceptor neurons. J Neurosci 33, 16033–16044.

Wang, S., Shi, Y., Shu, S., Guyenet, P.G., and Bayliss, D.A. (2013b). Phox2b-expressing retrotrapezoid neurons are intrinsically responsive to H+ and CO2. J Neurosci 33, 7756–7761.

Wang, W., Bradley, S.R., and Richerson, G.B. (2002). Quantification of the response of rat medullary raphe neurones to independent changes in pH(o) and P(CO2). J Physiol 540, 951–970.

Wang, W., Pizzonia, J.H., and Richerson, G.B. (1998). Chemosensitivity of rat medullary raphe neurones in primary tissue culture. J Physiol 511, 433–450.

Wang, W., Tiwari, J.K., Bradley, S.R., Zaykin, R.V., and Richerson, G.B. (2001). Acidosis-stimulated neurons of the medullary raphe are serotonergic. J Neurophysiol 85, 2224–2235.

Wenker, I.C., Sobrinho, C.R., Takakura, A.C., Moreira, T.S., and Mulkey, D.K. (2012). Regulation of ventral surface CO2/H+-sensitive neurons by purinergic signalling. J Physiol 590, 2137–2150.

West, J.B. (2012). Respiratory Physiology: the Essentials. (Philadelphia: Wolters Kluwer Health, Lippincott Williams & Wilkins).

Wu, Y., Proch, K.L., Teran, F.A., Lechtenberg, R.J., Kothari, H., and Richerson, G.B. (2019). Chemosensitivity of Phox2b-expressing retrotrapezoid neurons is mediated in part by input from 5-HT neurons. J Physiol 597, 2741–2766.

Yu, J., Bippes, C.A., Hand, G.M., Muller, D.J., and Sosinsky, G.E. (2007). Aminosulfonate modulated pH-induced conformational changes in connexin26 hemichannels. J Biol Chem 282, 8895–8904.

