## Supplementary Material for "Connexin26 mediates CO_2_-dependent regulation of breathing via glial cells of the medulla oblongata"

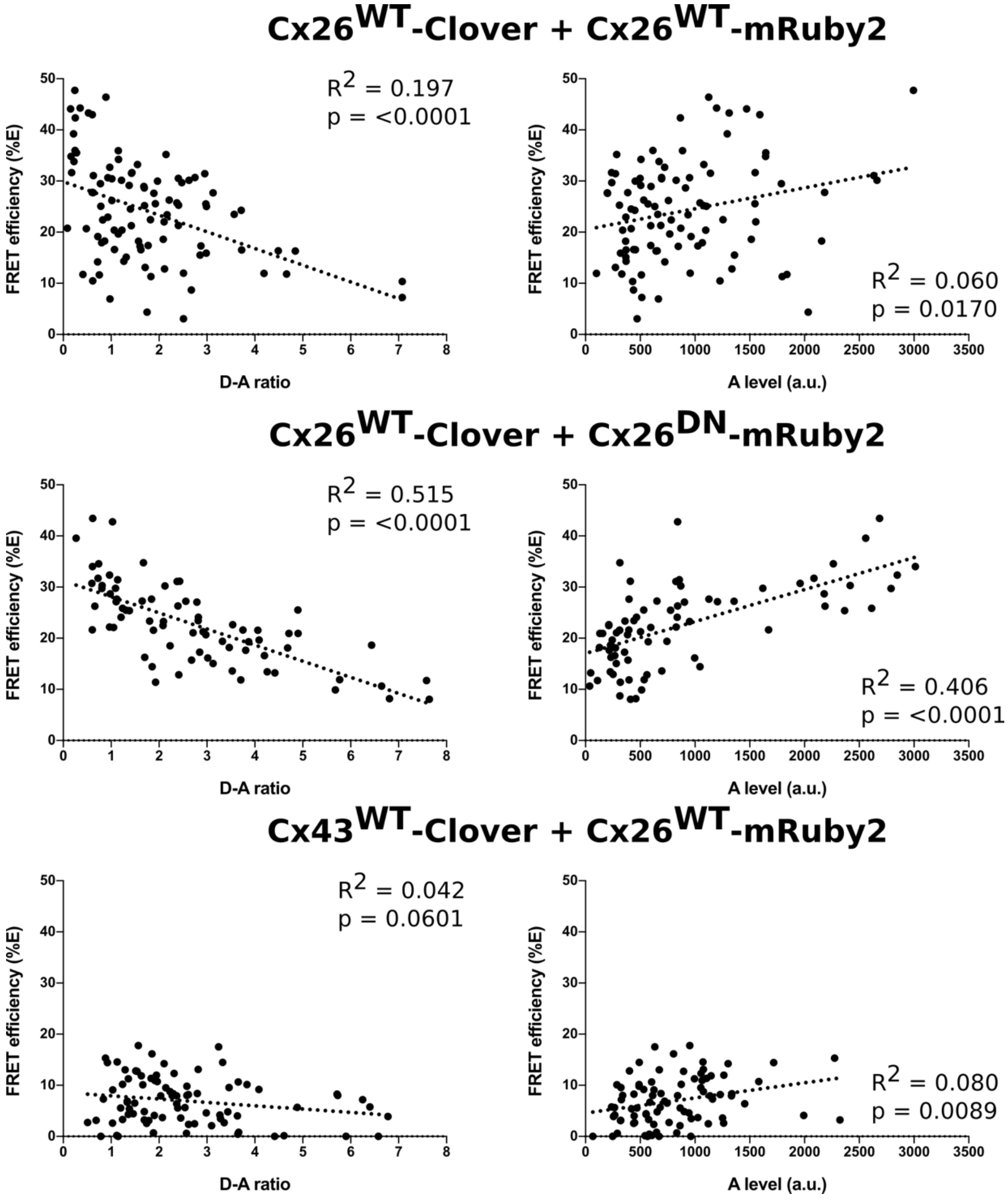

**Supplementary Figure 1.** The dependence of FRET efficiency (%E) on donor to acceptor (D-A) ratio -see Methods for calculations. A negative correlation between %E and D-A ratio is evidence for coassembly of the subunits into a hexamer (broadly, as the proportion of donor subunits grows relative to the acceptor subunits in the hexamer, the acceptor will saturate and FRET efficiency will decline). A positive correlation between %E and the amount of acceptor (A level), indicates random association between homomeric subunits in the membrane.

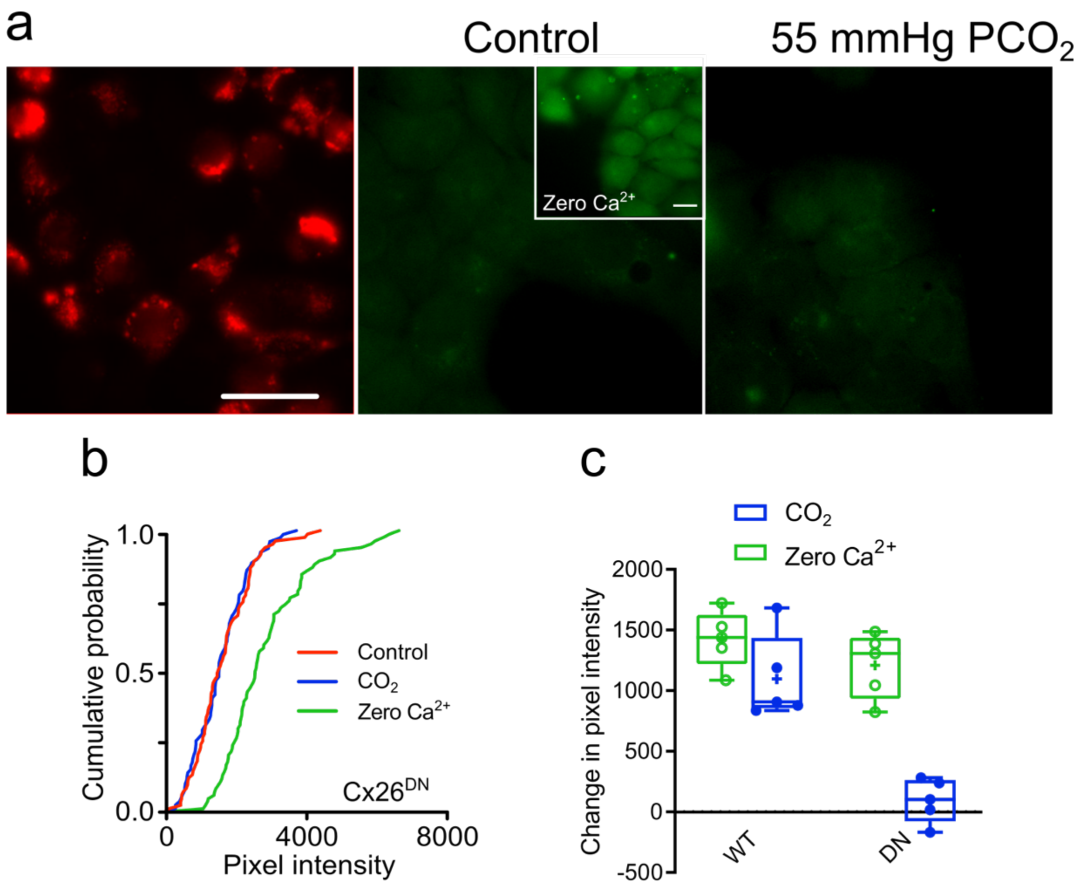

**Supplementary Figure 2.** Cx26[DN] is insensitive to $CO_2$. a) Images of mCherry expression, and dye loading in control and hypercapnic solutions. The inset demonstrates dye loading in response to zero $Ca^{2+}$ solutions. This will open hemichannels, and is a positive control to demonstrate the existence of function hemichannels. b) Cumulative probability histograms for dye loading in Cx26[DN]-expressing HeLa cells. There is no effect of $CO_2$ (55 mmHg stimulus) but zero $Ca^{2+}$ causes a rightward shift in pixel intensity. c) Median change in median pixel intensity for Cx26[WT] and Cx26[DN], showing that the wild type hemichannels responds to $CO_2$ but Cx26[DN] does not.

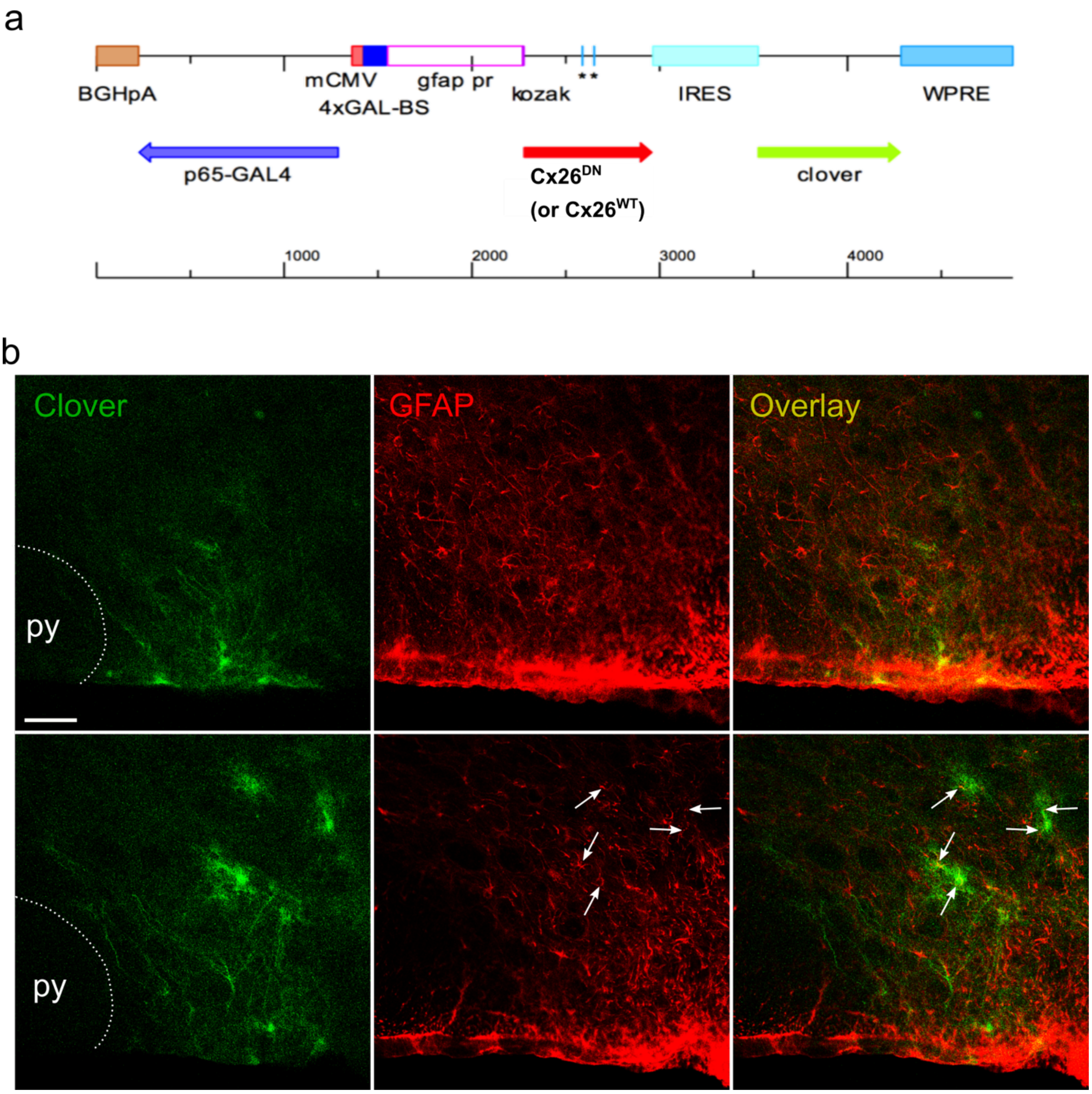

**Supplementary Figure 3.** a) Lentiviral construct sequences for *in vivo* expression of $Cx26^{WT}$ or $Cx26^{DN}$. b) Demonstration of selective expression of $Cx26^{DN}$ in GFAP+ cells in coronal sections of the caudal parapyramidal area. Montage of Clover fluorescence (green) and GFAP immunoreactivity (red) with overlay. Each image is a single optical section obtained by confocal microscopy. The top row shows ventral cells (within 50 µm of ventral surface). 42/42 Clover labelled cells in this position displayed GFAP immunoreactivity. The bottom row shows more dorsal cells, all of which had typical astrocytic morphology. The white arrows point to puncta of GFAP immunoreactivity present in these cells. Overall 39/40 Clover-labelled cells in the dorsal position had GFAP immunoreactivity. Scale bar 50 µm; py, pyramids.

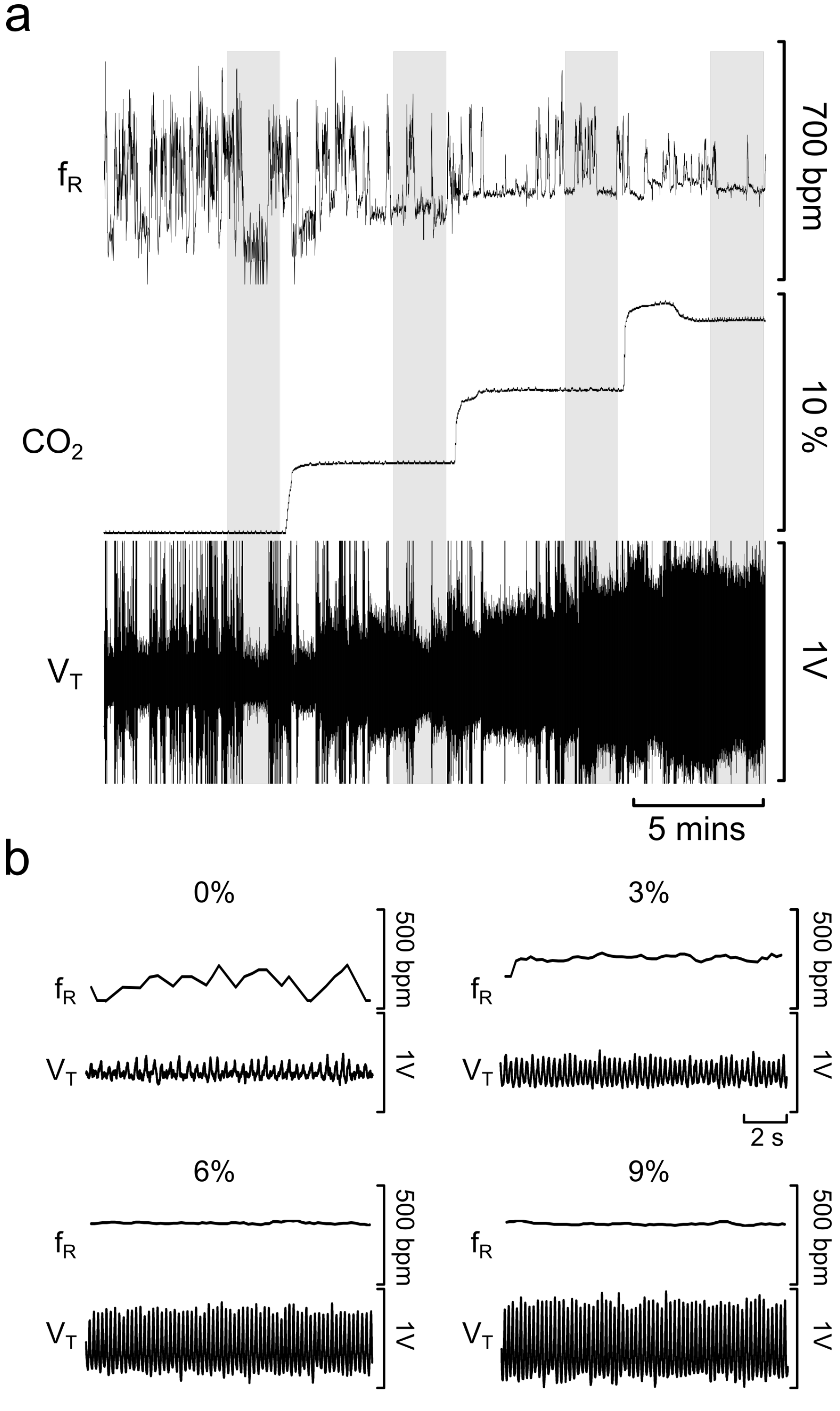

**Supplementary Figure 4.** Raw plethysmography data. The grey regions show those parts of the experiment selected for analysis of respiratory frequency ($f_R$) and respiratory flow traces ($V_T$) for calculation of tidal volume. Large excursions on the $V_T$ trace are movement artefacts.

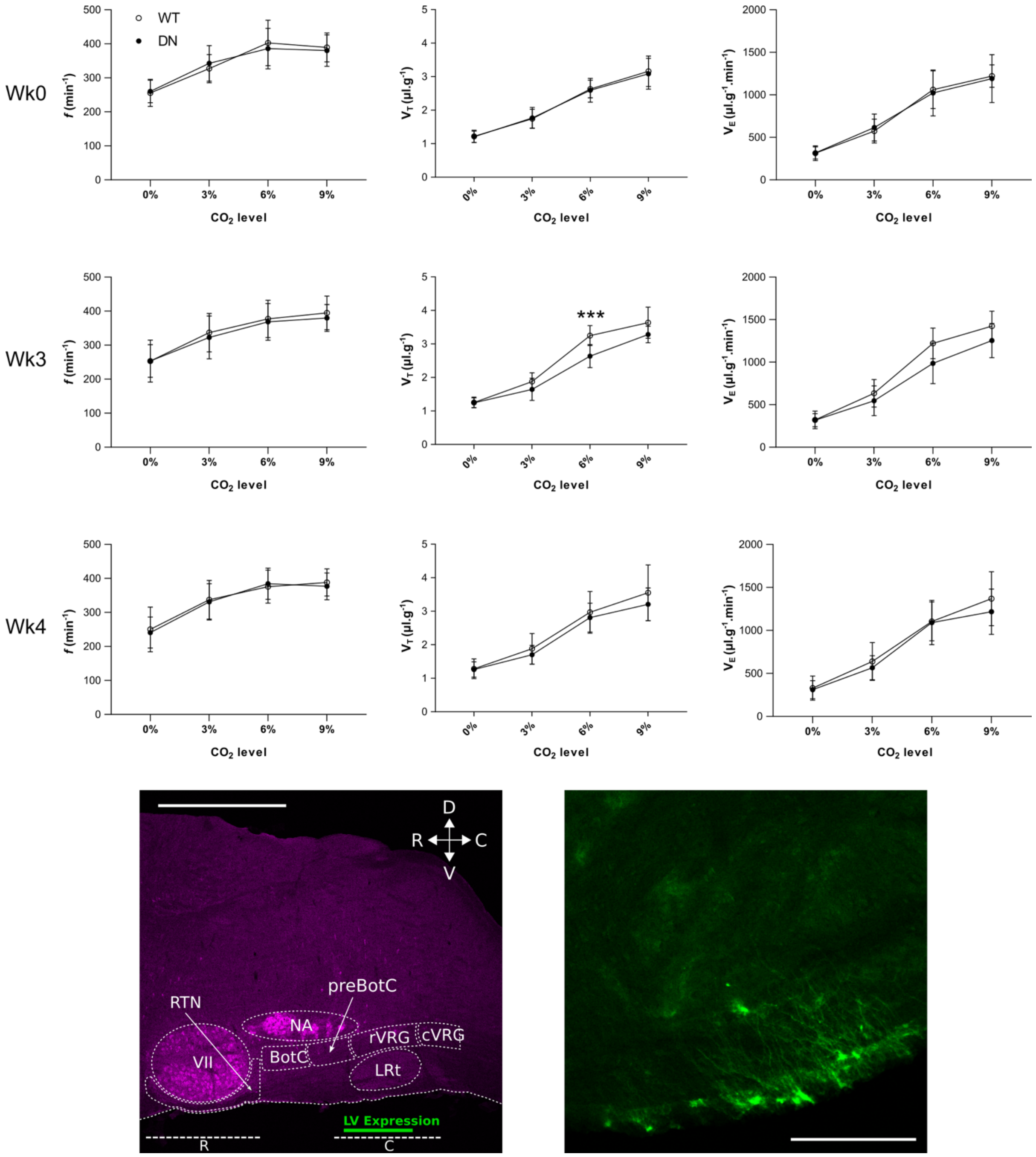

**Supplementary Figure 5.** Pilot study showing that $Cx26^{DN}$ expression in caudal area near the lateral reticular nucleus (LRt) reduced the tidal volume and minute ventilation responses to 6% $CO_2$ compared to $Cx26^{WT}$. This effect was apparent at week 3 after viral transduction but had disappeared at week 4. Data plotted as mean ± 95% confidence limits n=6 for $Cx26^{WT}$, n=8 for $Cx26^{DN}$. Scale bars 1mm (left), 200 µm (right). For $V_T$ at 3 weeks the 2-way mixed effect ANOVA was significant p = 0.019 and post-hoc t-tests (one-sided) showed that $V_T$ was reduced at 6% $CO_2$ (p=0.005, ***); and a marginal effect at 9% $CO_2$ (p=0.049, which did not pass the false discovery criterion). The 2-way mixed effects ANOVA showed that the differences for minute ventilation ($V_E$) were not significant (p=0.091).

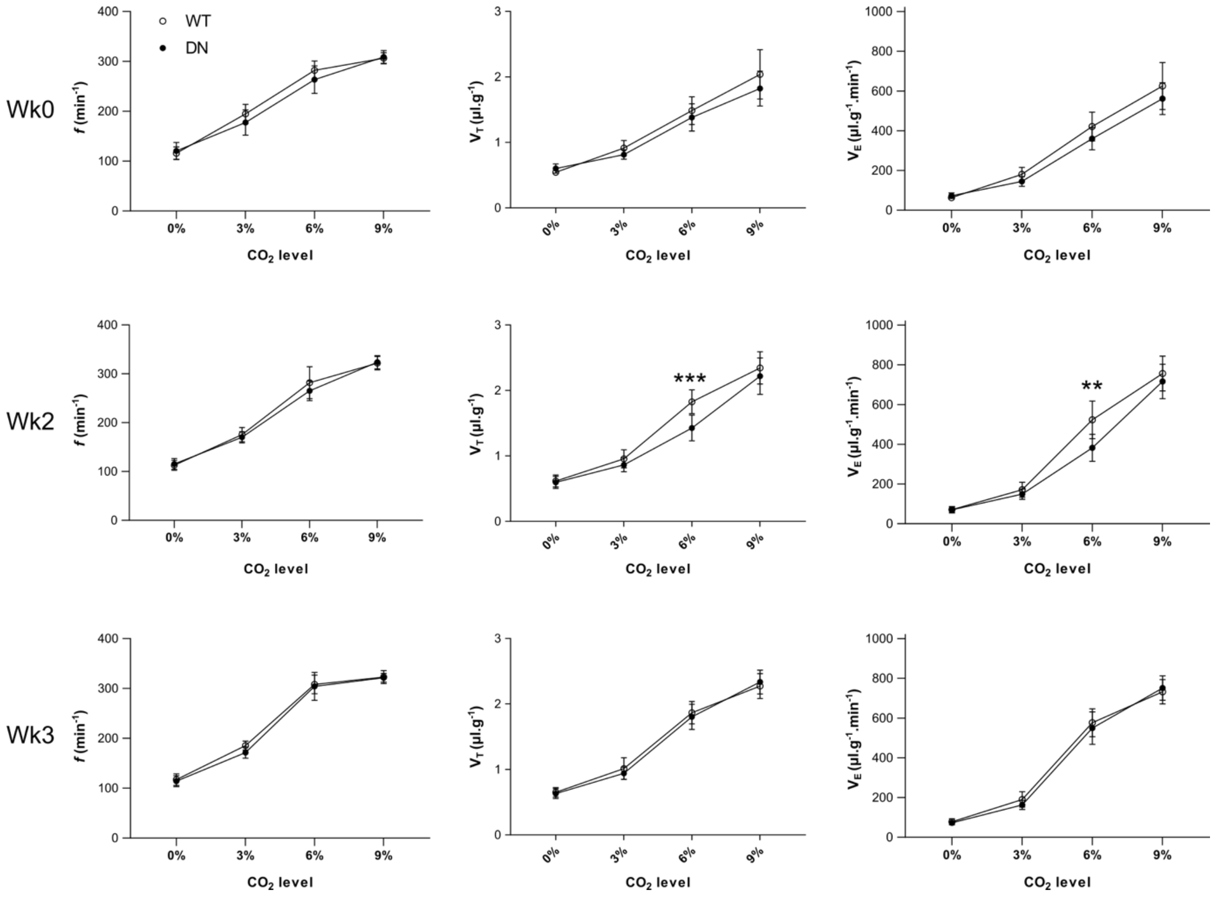

**Supplementary Figure 6.** Data from Figure 5 of main text plotted as mean ± 95% confidence limits. ***, $p=0.002$; **, $p=0.007$, post hoc comparison after 2-way mixed effects ANOVA.

**Supplementary Table 1**

| | Cx26^WT^ $\Delta V_T$ (µl.g$^{-1}$) | | | Cx26^DN^ $\Delta V_T$ (µl.g$^{-1}$) | | |
|---|---|---|---|---|---|---|
| | 3% | 6% | 9% | 3% | 6% | 9% |
| **Mean** | **0.34** | **1.21** | **1.73** | **0.27** | **0.83** | **1.62** |
| SD | 0.14 | 0.24 | 0.33 | 0.11 | 0.26 | 0.32 |
| **Median** | **0.34** | **1.25** | **1.73** | **0.28** | **0.82** | **1.46** |
| LQ, UQ | 0.29, 0.43 | 1.05,1.33 | 1.57, 1.95 | 0.19, 0.36 | 0.66, 0.89 | 1.38, 1.88 |

| % reduction by Cx26$^{DN}$ | 3% | **6%** | 9% |
|---|---|---|---|
| Means | 21 | **31** | 6 |
| Medians | 18 | **35** | 16 |

Data from Figure 5, and Supplementary Figure 6 – to quantify the effect on the change in tidal volume ($\Delta V_T$) evoked by 3, 6 and 9% inspired $CO_2$ of expression of Cx26$^{DN}$ relative to expression of Cx26$^{WT}$ in the caudal parapyramidal area. Cx26$^{DN}$ caused a reduction of $\Delta V_T$ of 31% (from means) and 35% (from medians). Although there were reductions $\Delta V_T$ at 3 and 9% these were not statistically significant.

**Supplementary Table 2**

| | Cx26$^{WT}$ $\Delta V_E$ (µl.g$^{-1}$.min$^{-1}$) | | | Cx26$^{DN}$ $\Delta V_E$ (µl.g$^{-1}$.min$^{-1}$) | | |
|---|---|---|---|---|---|---|
| | 3% | 6% | 9% | 3% | 6% | 9% |
| **Mean** | **101** | **452** | **685** | **78** | **312** | **646** |
| SD | 42 | 143 | 125 | 33 | 100 | 117 |
| **Median** | **96** | **465** | **690** | **73** | **309** | **648** |
| LQ, UQ | 83,129 | 404, 551 | 616, 773 | 52,108 | 255, 349 | 549, 728 |

| % reduction by Cx26$^{DN}$ | 3% | **6%** | 9% |
|---|---|---|---|
| Means | 22 | **31** | 6 |
| Medians | 23 | **34** | 6 |

Data from Figure 5, and Supplementary Figure 6 – to quantify the effect on the change in minute ventilation ($\Delta V_E$) evoked by 3, 6 and 9% inspired $CO_2$ of expression of Cx26$^{DN}$ relative to expression of Cx26$^{WT}$ in the caudal parapyramidal area. Cx26$^{DN}$ caused a reduction of $\Delta V_E$ of 31% (from means) and 34% (from medians). Although there were reductions of $\Delta V_E$ at 3 and 9% these were not statistically significant.

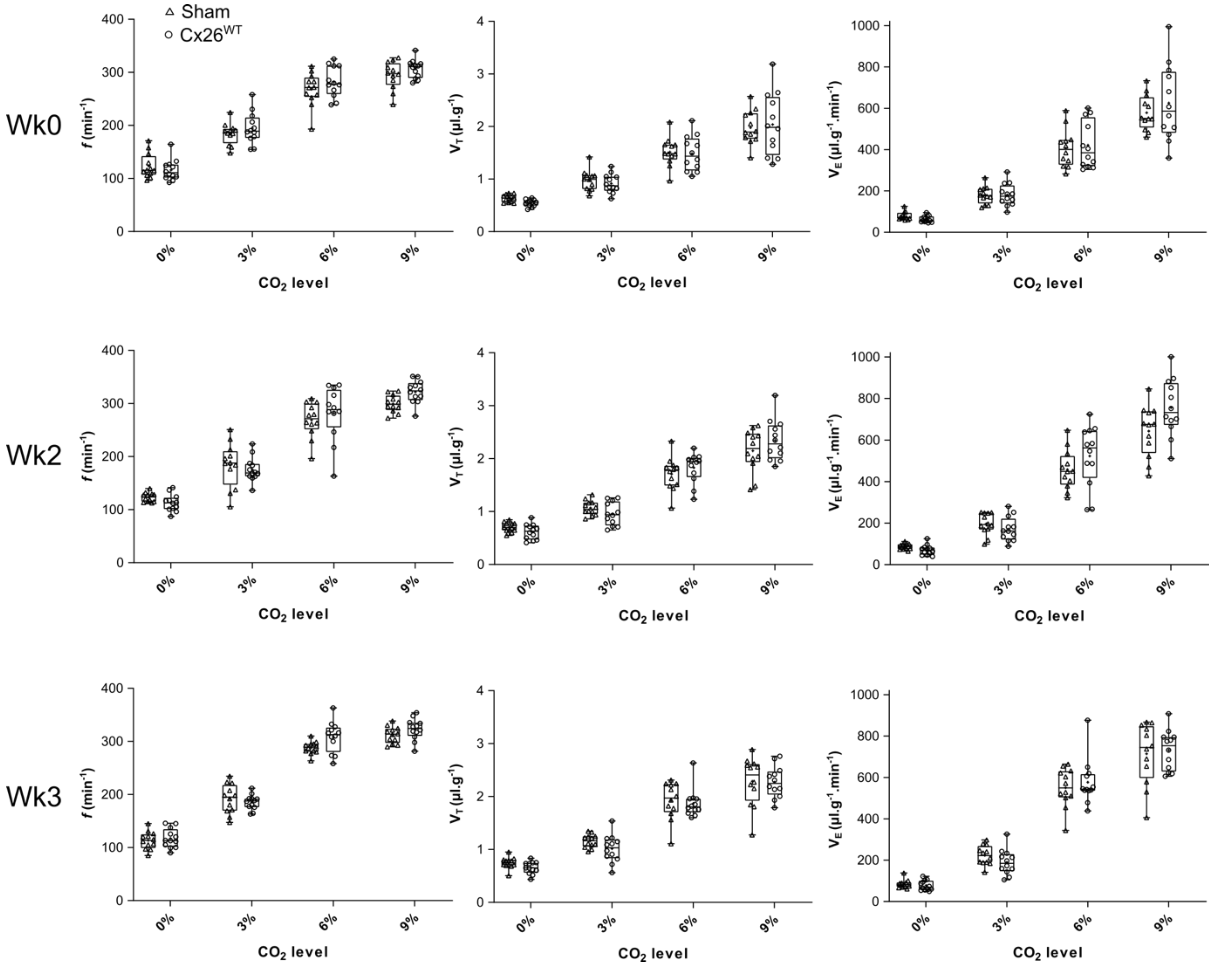

**Supplementary Figure 7.** Overexpression of Cx26$^{WT}$ does not increase sensitivity to $CO_2$ compared to sham operated mice. Cx26$^{WT}$ data same as in Figure 5 of main text, the sham operation was performed as part of same study as that of Figure 5.

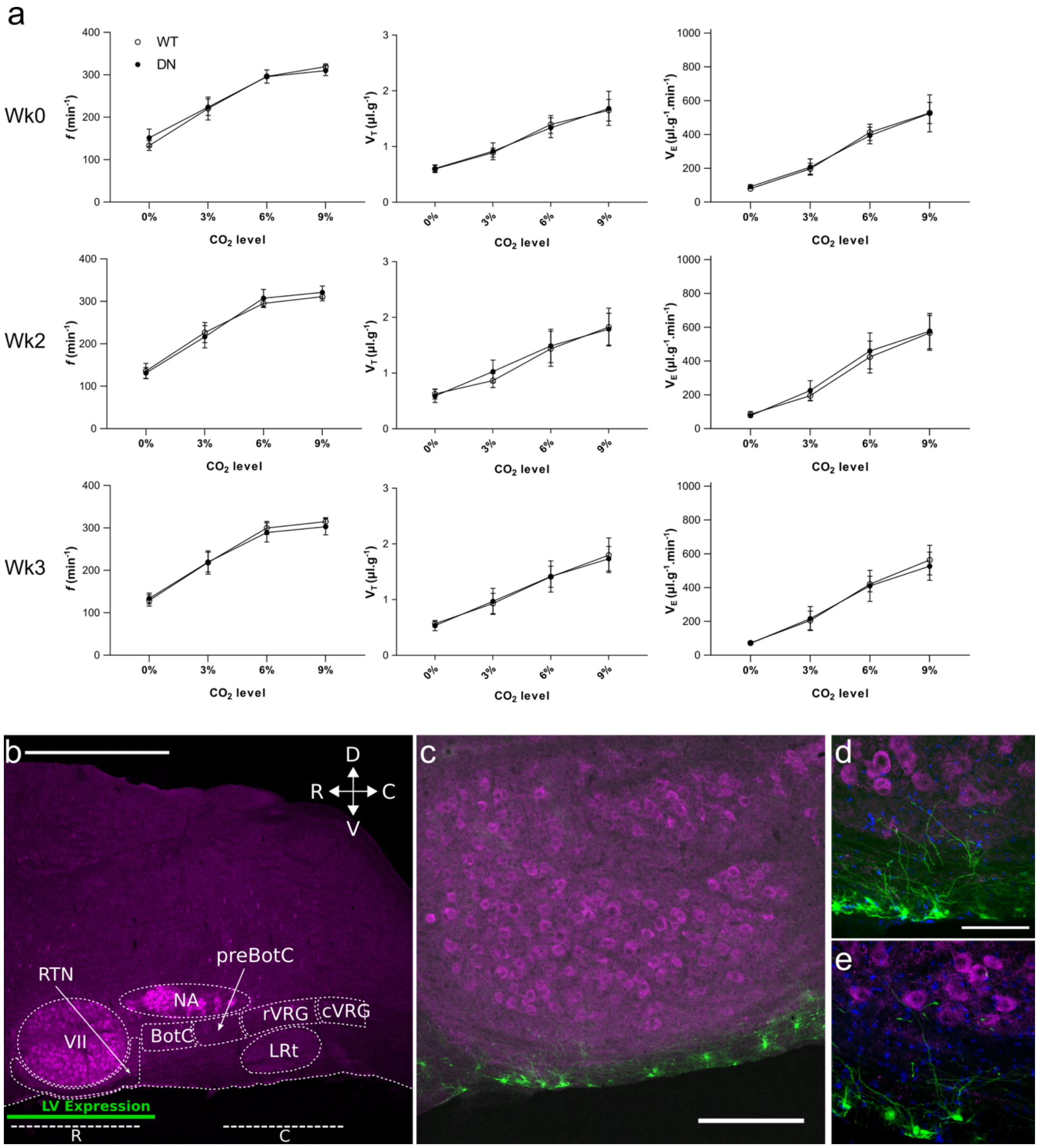

**Supplementary Figure 8.** a) There is no effect of $Cx26^{DN}$ on the $CO_2$ sensitivity of beathing when expressed in glial cells of the RTN. Data plotted as mean ± 95% confidence limits. n=8 for both $Cx26^{WT}$ and $Cx26^{DN}$. b) Location of LV expression. c-e) Micrographs of RTN glial cells transduced. Scale bars: b) 1 mm; c) 200 µm; d,e) 50 µm.

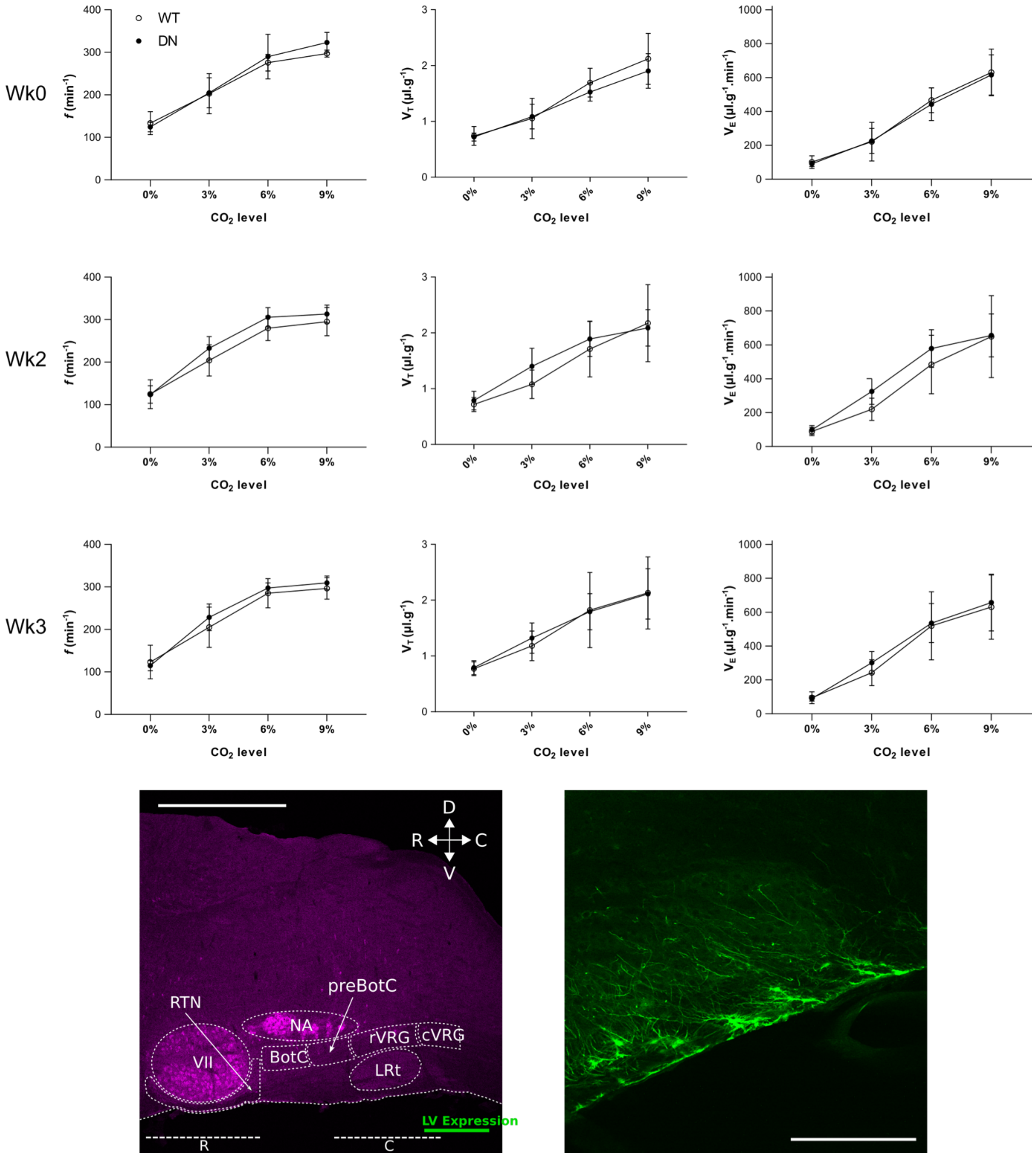

**Supplementary Figure 9.** Expression of Cx26[DN] in the very caudal medulla oblongata has no effect on the $CO_2$-sensitivity of breathing. n=6 for Cx26[WT] and Cx26[DN]. Scale bars: left) 1 mm; right) 200 µm.

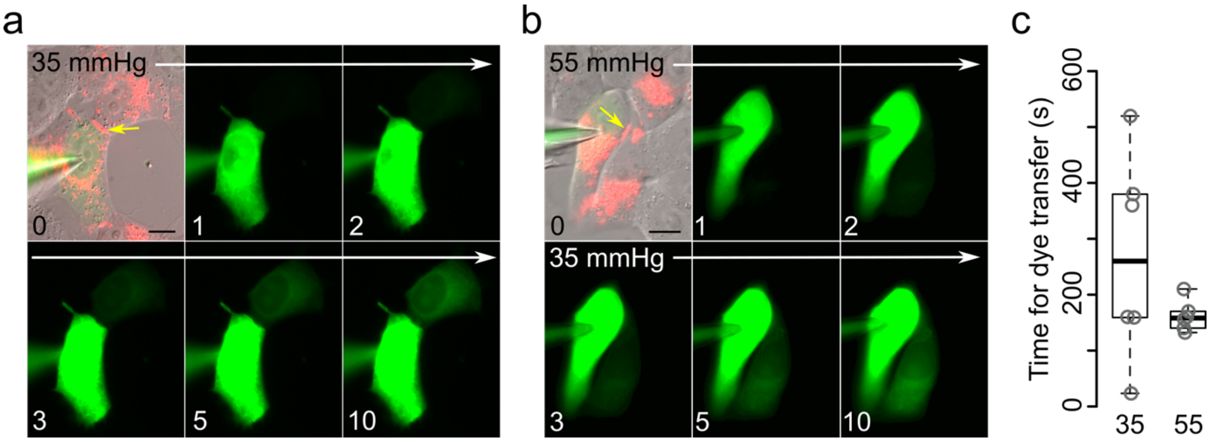

**Supplementary Figure 10. Cx26$^{DN}$ forms functional gap junctions that are insensitive to $CO_2$.** Recordings were made with patch electrodes from coupled HeLa cells expressing Cx26$^{DN}$ tagged with mCherry. A patch pipette was used to introduce 2-Deoxy-2-[(7-nitro-2,1,3-benzoxadiazol-4-yl)amino]-D-glucose (NBDG) -a fluorescent glucose analogue -into a single cell (the *donor*) of a coupled pair. The time taken for the dye to diffuse into a coupled cell and achieve 10% of the fluorescence intensity of the donor cell was determined.

a) Montage of images showing superimposed DIC, mCherry (tagged to Cx26$^{DN}$, red) and NBDG at 0, 1, 2, 3, 5 and 10 minutes after breakthrough at a $PCO_2$ of 35 mmHg - fluorescence was observed in the coupled cell 3 minutes after breakthrough. Yellow arrows indicate gap junction plaque between cells, scale bar 20 µm.

b) Montage of images showing superimposed DIC, mCherry (tagged to Cx26$^{DN}$) and NBDG at 0, 1, 2, 3, 5 and 10 minutes after breakthrough starting at a $PCO_2$ of 55 mmHg, and transferring to 35 mmHg after 2 minutes. The presence of high $PCO_2$ does not delay entry of NBDG - fluorescence in the coupled cell was observed 3 minutes after breakthrough. Yellow arrows indicate gap junction plaque between cells, scale bar 20 µm.

c) Plot of time taken for fluorescence of coupled cell to reach 10% of that of the donor. The starting $PCO_2$ makes no difference to the time for permeation of NBDG indicating that the gap junction formed by Cx26$^{DN}$ is insensitive to $CO_2$ (n=6 for each condition).
